# Rapid degradation of Histone Deacetylase 1 (HDAC1) reveals essential roles in both gene repression and active transcription

**DOI:** 10.1101/2024.06.19.599716

**Authors:** David M English, Samuel N Lee, Khadija A Sabat, India M Baker, Trong Khoa Pham, Mark O Collins, Shaun M Cowley

**Affiliations:** Department of Molecular and Cell Biology, Henry Wellcome Building, University of Leicester, Leicester, LE1 7RH, United Kingdom; Cambridge Stem Cell Institute & Department of Haematology, Jeffrey Cheah Biomedical Centre, Cambridge Biomedical Campus, University of Cambridge, Puddicombe Way, Cambridge, CB2 0AW, United Kingdom; School of Biosciences, University of Sheffield, Sheffield S10 2TN, UK; biOMICS Mass Spectrometry Facility, University of Sheffield, Sheffield S10 2TN, UK

## Abstract

Histone Deacetylase 1 (HDAC1) removes acetyl groups from lysine residues on the core histones, a critical step in the regulation of chromatin accessibility. Despite histone deacetylation being an apparently repressive activity, suppression of HDACs causes both up- and down-regulation of gene expression. Here we exploited the degradation tag (dTAG) system to rapidly degrade HDAC1 in embryonic stem cells (ESCs) lacking its paralog, HDAC2. Unlike HDAC inhibitors that lack isoform specificity, the dTAG system allowed specific degradation and removal of HDAC1 in <1 hour (100x faster than genetic knockouts). This rapid degradation caused increased histone acetylation in as little as 2 hours, with H2BK5 and H2BK11 being the most sensitive. The majority of differentially expressed genes following 2 hours of HDAC1 degradation were upregulated (275 genes up vs 15 down) with increased proportions of downregulated genes observed at 6 (1,153 up vs 443 down) and 24 hours (1,146 up vs 967 down) respectively. Upregulated genes showed increased H2BK5ac and H3K27ac around their transcriptional start site (TSS). In contrast, decreased acetylation of super-enhancers (SEs) was linked to the most strongly downregulated genes. These findings suggest a paradoxical role for HDAC1 in the maintenance of histone acetylation levels at critical enhancer regions required for the pluripotency-associated gene network.

## Introduction

Acetylation of the ε-amino group of the lysine side chain is a post-translational modification that occurs on thousands of proteins. However, the bulk (74%) of this acetylation is found on histone proteins with their lysine-rich tails (1, 2). Lysine acetylation on the N-terminal tails of histone proteins is an essential mechanism for the regulation of chromatin structure and gene expression (3). The addition of an acetyl group masks the positive charge of the lysine residue, loosens the interactions of the histones with DNA and reduces interactions between neighbouring nucleosomes. Additionally, the acetyl lysine acts as a binding site for proteins with bromodomains, that are found in general transcription factors and chromatin remodelling complexes (3-5). The addition of the acetyl group can occur via the action of histone acetyltransferases (HATs) or through chemical reaction with acetyl-CoA, which has particularly been noted in the favourable conditions of the mitochondria (6, 7). The family of histone deacetylases (HDACs) are responsible for removing the acetyl group from lysine residues, with class I HDACs (HDAC1/2/3/8) of particular importance for gene regulation (8-11).

HDAC1 and HDAC2 (HDAC1/2) are sister enzymes (83% sequence similarity) that are recruited into six distinct co-repressor complexes: SIN3 (12, 13), NuRD (14), CoREST (15), MiDAC (16), MIER (17, 18) and RERE (19). With a couple of notable exceptions, *Hdac1* and *Hdac2* are for the most part functionally redundant in most tissue types, producing only mild phenotypes (summarized in (20)). In contrast, double knockout (DKO) of *Hdac1/2* produces severe phenotypes, which is unsurprising given that HDAC1/2 are responsible for over half of the total deacetylase activity in cells (9, 21). The use of both genetic knockouts (KOs) and class I HDAC specific inhibitors has highlighted the importance of HDAC1/2 in removing acetylation from high stoichiometry sites on histone tails (1, 9, 21). Despite the removal of acetyl marks appearing to be a repressive function, ChIP-seq revealed that HDAC1/2 and their associated complexes are found located at sites of active transcription (22, 23). In addition, transcriptome studies have shown that the loss of HDAC1/2 results in equal numbers of genes with increased and decreased expression, highlighting a role for histone deacetylation in active gene expression (9).

Studies utilising HDAC inhibitors and conditional knockouts have generated data that has proven very useful, but they have been limited by either a lack of isoform specificity or the very stable nature of the HDAC1/2 complexes themselves (half-life >24 hours) (9, 24-26). To bypass this, we have created a cell line that exploits the degradation tag (dTAG) system (27), to rapidly degrade HDAC1 (in the absence of HDAC2). The lethality of HDAC1/2 removal which took 4-5 days to become apparent in conditional DKO cells (9) is recapitulated here in just 24 hours. Substantial increases in acetylation were detected on key histone sites within 2 hours of dTAG treatment. These increases in acetylation were accompanied by hundreds of transcriptional changes in just 2 hours with numbers stretching into the thousands only 6 hours post-dTAG^V-^1 (28) treatment. Combined analysis of these transcriptional changes with locus-specific changes in acetylation determined by ChIP-seq helped to reveal how changes in acetylation are linked to gene expression and active gene transcription.

## Methods

### Cell culture

All experiments described here used *Hdac1/2^lox/lox^* cells (9) that were engineered to express an HDAC1-FKBP12^F36V^-Flag construct (referred to as HDAC1-FKBP cells) using Lipofectamine 2000 as detailed below. Cells were cultured in M15+LIF media as described previously (9). To induce deletion of endogenous *Hdac1/2* cells were treated with 1 μM 4-hydroxytamoxifen (OHT) for 4 days before use in experiments.

### Lipofectamine 2000 transfection

The HDAC1-FKBP cell line used in this study was created using Lipofectamine 2000 (Invitrogen) stable transfection of *Hdac1/2^lox/lox^* ESCs created previously (9). Transfections were carried out as described previously (29), with the PROTEX service at the University of Leicester cloning the HDAC1-FKBP12^F36V^-Flag construct into a *piggyBac* insertion vector.

### dTAG treatments

Cells in which endogenous *Hdac1/2* had been knocked out (as described above) were treated with 50 nM dTAG-13 or 100 nM dTAG^V^-1 (as stated in figure legends) or an equivalent amount of DMSO as a solvent control. Cells were then harvested using an appropriate volume of TrypLE (Gibco) for the size of culture plate following two washes with PBS to remove dead cells (dead cells were kept for viability studies) and used for the downstream experiments described below. Successful removal of endogenous HDAC1/2 and degradation of the HDAC1-FKBP12^F36V^ proteins were confirmed by western blotting with HDAC1/2 antibodies.

### Cell viability

To capture all cells (both adherent and floating) cell culture media was retained and transferred to a 15 mL tube. Cells were then detached with TrypLE that was neutralised using M15 media, this cell suspension was then transferred into the same 15 mL tube as the dead cells and the suspension mixed. Cell counts were performed using Trypan Blue (Bio-Rad) and a Bio-Rad TC20 automated cell counter following the manufacturer’s instructions. Unpaired t-tests were then conducted using graphpad prism to determine if there were any significant differences in viability based on the HDAC1/2 status of the cells.

### Propidium iodide cell cycle analysis

250,000 HDAC1-FKBP cells were seeded in 6 well plates 24 hours prior to treatment with 100 nM dTAG^V^-1 for 2, 6 or 24 hours or DMSO for 24 hours. Cells were then harvested and used for cell cycle distribution analysis using propidium iodide staining as described in (10). The gating strategy used is shown in Supplementary Fig S1. A full breakdown of propidium iodide FACS results are shown in Supplementary table S1.

### Western blotting

Western blots were performed as described previously (9). Primary antibodies used to probe membranes are shown in the table below:

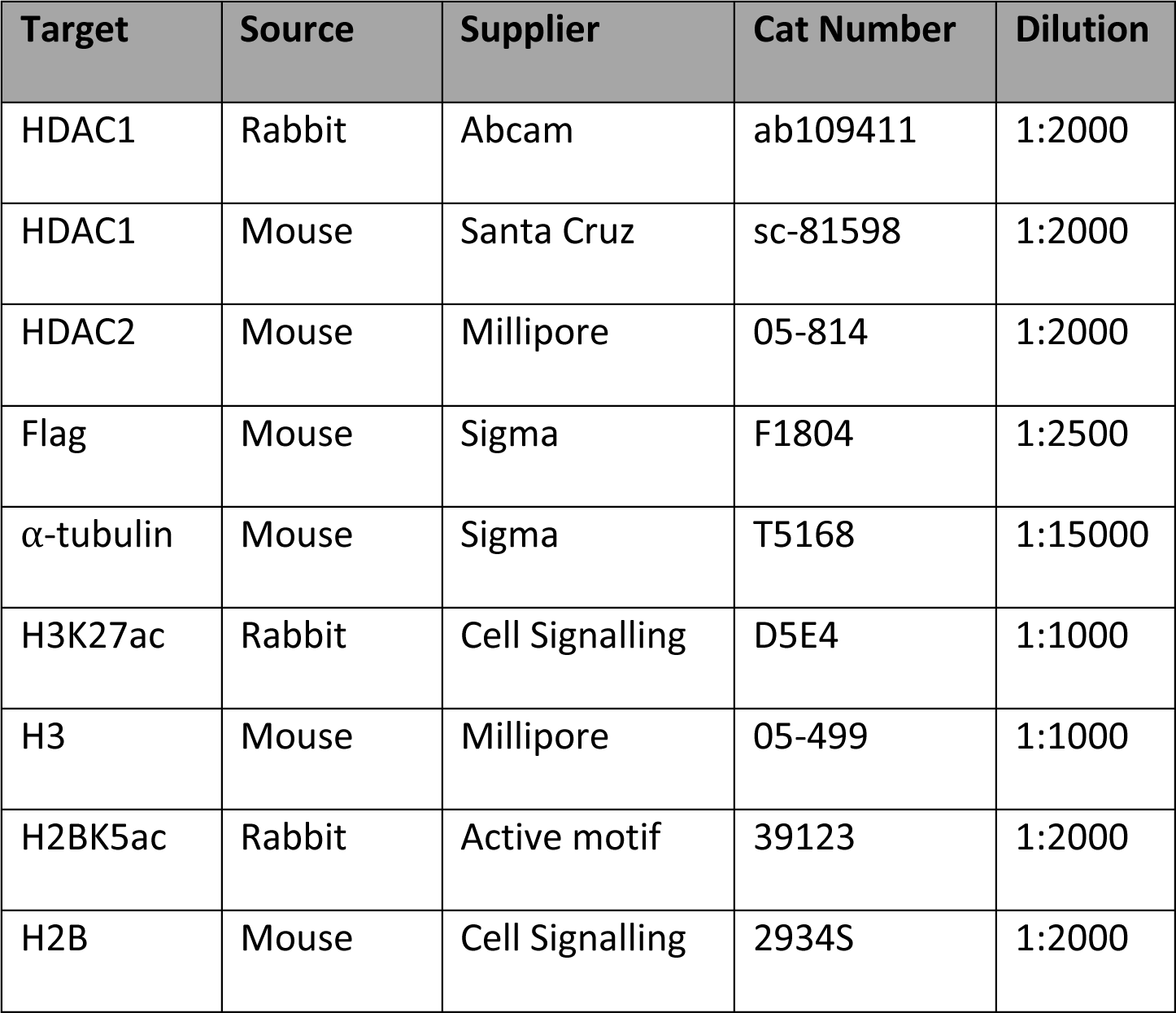

| **Target** | **Source** | **Supplier** | **Cat Number** | **Dilution** |
| --- | --- | --- | --- | --- |
| HDAC1 | Rabbit | Abcam | ab109411 | 1:2000 |
| HDAC1 | Mouse | Santa Cruz | sc-81598 | 1:2000 |
| HDAC2 | Mouse | Millipore | 05-814 | 1:2000 |
| Flag | Mouse | Sigma | F1804 | 1:2500 |
| α-tubulin | Mouse | Sigma | T5168 | 1:15000 |
| H3K27ac | Rabbit | Cell Signalling | D5E4 | 1:1000 |
| H3 | Mouse | Millipore | 05-499 | 1:1000 |
| H2BK5ac | Rabbit | Active motif | 39123 | 1:2000 |
| H2B | Mouse | Cell Signalling | 2934S | 1:2000 |

### Mass spectrometry analysis of histone acetylation

To obtain histone proteins a whole cell extract was first made by lysing cell pellets from confluent 10 cm plates in NP-40 lysis buffer (50 mM Tris-HCl pH 8, 150 mM NaCl, 1 mM EDTA, 1% NP-40, 10% glycerol, protease inhibitor cocktail (Sigma; P8340)) for 30 minutes. Lysates were cleared by centrifugation at 14,000 rpm at 4 °C for 15 minutes. The supernatant containing the whole cell extract was transferred to a fresh 1.5 mL tube and histones were extracted from the remaining pellet by overnight incubation (20 hours) in 50 μL of 0.4 N H_2_SO_4_. Samples were again centrifuged at 14,000 rpm at 4 °C for 15 minutes, and the supernatant containing the histone proteins was transferred to a fresh 1.5 mL tube. The samples were then neutralised using 0.8 N NaOH. Histones were derivatised according to (30). Briefly, 10 mg of histone samples in 50 mL of 50 mM ammonium bicarbonate were incubated with 16.6 mL of propionylation reagent (1:3 v/v propionic anhydride in acetonitrile) for 15 minutes at 37 °C with agitation in a thermomixer at 900 rpm. Samples were dried down in a vacuum concentrator and the derivatisation was repeated. 10 mg of derivatised and underivatised histone samples were digested in 50 mL of 50 mM ammonium bicarbonate with 1 mg of trypsin (Pierce, sequencing grade) for 2 hours at 37 °C with agitation in a thermomixer at 900 rpm. Digests were desalted using C18 spin columns (Pierce) according to the manufacturer’s protocol. Eluted peptides were dried in a vacuum concentrator and resuspended in 0.5% formic acid for LC-MS/MS analysis. Each sample was analysed using nanoflow LC-MS/MS using an Orbitrap Elite (Thermo Fisher) hybrid mass spectrometer equipped with a nanospray source, coupled to an Ultimate RSLCnano LC System (Dionex). Peptides were desalted online using a nano-trap column, 75 μm I.D.X 20mm (Thermo Fisher) and then separated using a 120-min gradient from 5 to 35% buffer B (0.5% formic acid in 80% acetonitrile) on an EASY-Spray column, 50 cm × 50 μm ID, PepMap C18, 2 μm particles, 100 Å pore size (Thermo Fisher). The Orbitrap Elite was operated in decision tree mode with a cycle of one MS (in the Orbitrap) acquired at a resolution of 120,000 at m/z 400, a scan range 375-1600, with the top 10 most abundant multiply charged (2+ and higher) ions in a given chromatographic window subjected to MS/MS fragmentation in the linear ion trap using CID and ETD depending on the charge state and m/z (the default decision tree settings were used). An FTMS target value of 1e6 and an ion trap MSn target value of 1e4 were used with the lock mass (445.120025) enabled. Maximum FTMS scan accumulation time of 200 ms and maximum ion trap MSn scan accumulation time of 50 ms were used. Dynamic exclusion was enabled with a repeat duration of 45 s with an exclusion list of 500 and an exclusion duration of 30 s.

### Mass spectrometry data analysis

Raw mass spectrometry data were analysed using MaxQuant version 1.6.2.6 (31). The following parameters were used to search against a mouse proteome: digestion set to Trypsin/P with 3 missed cleavages, methionine oxidation, N-terminal protein acetylation and lysine acetylation set as the variable modifications. Additionally, propionylation was set as a variable modification for derivatised samples with the number of missed cleavages set to 5. A protein and peptide FDR of 0.01 were used for identification level cut-offs based on a decoy searching database strategy. Mass spectrometry data is available in Supplementary table S2. The mass spectrometry proteomics data have been deposited to the ProteomeXchange Consortium via the PRIDE (32) partner repository with the dataset identifier PXD053032.

### RNA-seq

Total RNA was extracted using a Promega Maxwell 16 and quality checked using an Agilent Bioanalyzer by the NUCLEUS facility at the University of Leicester. mRNA library preparation and sequencing were carried out by Novogene (Cambridge, UK) with sequencing to a depth of 20 million reads using the NovaSeq 6000 PE150 platform. Bioinformatic analysis was performed as described in (10) with reads mapped to the mm10 genome index. The raw and processed files are stored at GEO SuperSeries (GSE269448). The bioconductor package TopGO was used for gene ontology (GO) analysis with the mouse genome wide annotation package org.Mm.eg.db.

### ChIP-seq

Native ChIP was performed as described in (33). 1 mL of diluted nucleosomes were incubated with 4 μL of anti-H3K27ac (Cell Signalling Technology D5E4) or 10 μL of anti-H2BK5ac (Active Motif 39123). Libraries were prepared for ChIP and matched input samples using the NEBNext Ultra II DNA library kit and indexed using NEBNext Multiplex Oligos. The NUCLEUS facility at the University of Leicester checked the average size and concentration of each library using an Agilent Bioanalyzer with a DNA high-sensitivity kit. Sequencing was conducted by Novogene (Cambridge, UK) with a sequencing depth of at least 25 million reads on the NovaSeq 6000 PE150 platform.

### ChIP-seq analysis

Raw reads were trimmed using trimmomatic before aligning to the mm10 genome sequence using Bowtie 2 using –no-mixed and –no-discordant options. SAM files were converted to BAM files, then sorted using samtools before duplicates were removed using sambamba. Peak calling was performed on ChIP (i.e. non-input) files using macs3 using default parameters. Diffbind was used to perform differential peak analysis. BAM files for individual replicates were merged using samtools merge. bigWigs were created from BAM files using bamCoverage (--binSize 20, --extendReads 150, --minMappingQuality 10, --normalizingUsing RPGC, --ignoreForNormalization chrX chrM, and mm10 blacklisted regions removed) (deepTools). Input files were subtracted from ChIP files using bigwigcompare (-binSize 20) (deepTools). bigWig files were uploaded to UCSC for viewing. Metaplots were created from bigWig files using deepTools programs computeMatrix to create a table of reads across defined regions (e.g. promoters, enhancers, TSS) and plotHeatmap and plotProfile were used to create the meta plots. The raw and processed files are stored in GEO SuperSeries (GSE269448).

For the analysis of promoter regions of upregulated genes, the coordinates for the regions from the TSS to -500 bp were downloaded from the UCSC genome table browser (34). This includes transcript variants and genes with multiple promoters giving 6202 promoter regions. For super-enhancer (SE) analysis we used the regions defined in (35) and for the promoter regions of SE-associated genes we downloaded the coordinates for the regions from the TSS to -500 bp from the UCSC genome table browser (34). For comparison between SE acetylation and the promoters of SE genes we removed promoter regions that overlapped SEs from our analysis. Violin plots were made using ggstatsplot, significance defined using a parametric t test with Holm-Bonferroni correction. The tracks for MED1 and p300 that are shown on Fig 6 were downloaded from NCBI GEO as referenced in (36, 37) and then viewed using the UCSC genome browser (38).

## Results

### Rapid HDAC1 degradation causes a loss of ESC viability and increased histone acetylation

Due to the long half-lives of HDAC1/2 (approximately 24 hours (26)) it takes at least 4 days for the complete loss of HDAC1/2 protein upon conditional knockout, blurring the distinction between primary and secondary effects (9). To overcome this, we stably expressed an HDAC1 protein with a C-terminal FKBP12^F36V^ tag to allow rapid degradation upon treatment with dTAG molecules (27). Initial treatment with 4-hydroxytamoxifen (OHT) removed endogenous HDAC1/2 protein, leaving only the HDAC1-FKBP12^F36V^ protein expressed after 5 days (Fig 1A, compare first two lanes. Upon treatment with dTAG-13 (or dTAG^V^-1 (Supplementary Fig S2)) the HDAC1-FKBP12^F36V^ protein is removed in just one hour (Fig 1A). Importantly, the functionality of the HDAC1-FKBP12^F36V^ protein is confirmed by a rescue of cell viability in the absence of endogenous HDAC1/2 (Fig 1B). This characteristic reduction in cell viability associated with HDAC1/2 loss (9) is observed as early as 24 hours after treatment with the dTAG molecule (Supplementary Fig S3). This is reflected in a 70% drop in viability (Fig 1B) and a proportional increase in the sub-G1 cell population following loss of HDAC1-FKBP12^F36V^ at 24 hours (Fig 1C).

**Figure 1.**
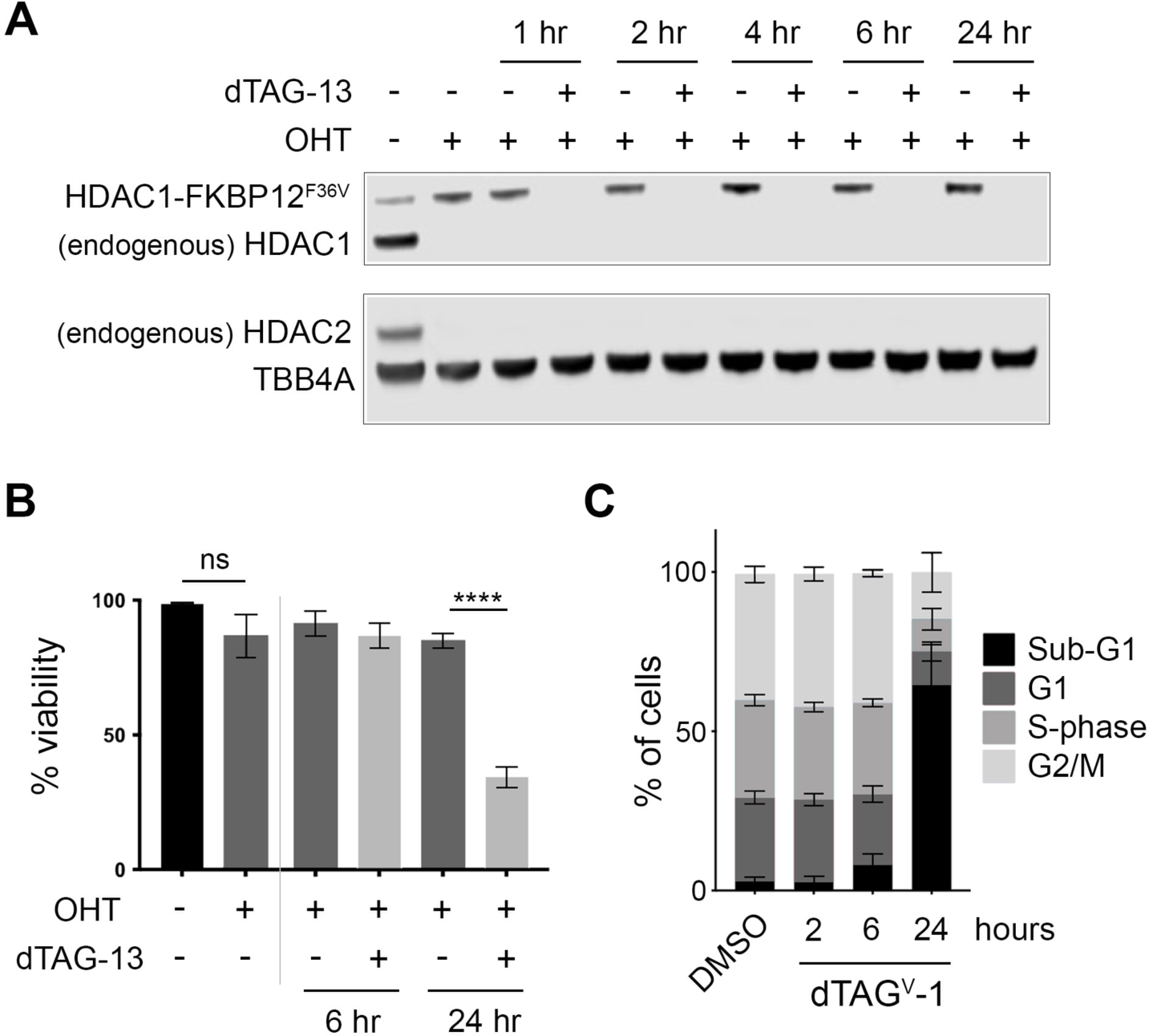
HDAC1-FKBP12^F36V^ is degraded rapidly by dTAG treatment, inducing cell death. (**A**) Western blot showing endogenous HDAC1/HDAC1-FKBP12^F36V^ (detected with ⍺-HDAC1), endogenous HDAC2 (detected with ⍺-HDAC2) when treated with OHT and 50 nM dTAG-13 as indicated, with ⍺-tubulin as a loading control. (**B**) Graph showing viability of HDAC1-FKBP cells following the indicated treatments with OHT and 50 nM dTAG-13 (n=3 -/+ SD) (ns = non-significant, **** P<0.0001). (**C**) Quantification of PI-FACS showing the percentage of HDAC1-FKBP cells in the indicated cell cycle stages following the indicated time of treatment with 100 nM dTAG^V^-1.

The acetylation of lysines within histone tails is highly dynamic with half-lives as short as 10 minutes for specific sites within H2B (39). The HDAC1-FKBP cells allowed us to examine histone substrates within 1 hour of degradation. To this end, we prepared histone extracts from cells at 2 and 6 hours of dTAG treatment. In initial western blots, we found significant increases to H3K27ac (approximately 2.2-fold) and H2BK5ac (approximately 1.8-fold) at 6 hours (Fig 2A). Both modifications are important markers of gene expression, with H3K27ac marking active enhancers (40) and H2BK5ac found at both enhancers and promoters (36). To explore the wider effects of rapid HDAC1 depletion on histone acetylation, we used mass-spectrometry at 2 and 6 hours post-dTAG treatment. We monitored increased acetylation levels at sites on H2B, H3 and H4 as well as macro-H2A (Fig 2B). The majority of sites responded strongly to HDAC1 removal, with only seven sites showing less than a 2-fold increase at 2 hours (macro-H2A.1K6, H3.1K14, H3.1K23, H3.1K27, H3.2K59, H3.3K27 and H4K16) and four at 6 hours (H3.1K14, H3.1K23, H3.2K59 and H4K16). At 6 hours many of the largest increases were found on histone H2B, particularly K5 and K11 (approximately 11-fold). On histone H3, H3K18 showed the greatest increase (approximately 4-fold), with K5 increasing the most on histone H4 (approximately 4.5-fold). Several of the sites we detected have previously been identified as sensitive to class I HDAC inhibitors, such as CI-994 (25), with H2BK20ac shown as an HDAC1/2 target in HDAC1/2 dTAG ESCs (36). These results show that nearly all sites of acetylation in histones are sensitive to the loss of HDAC1, with changes detected as early as 2 hours increasing progressively at 6 hours.

**Figure 2.**
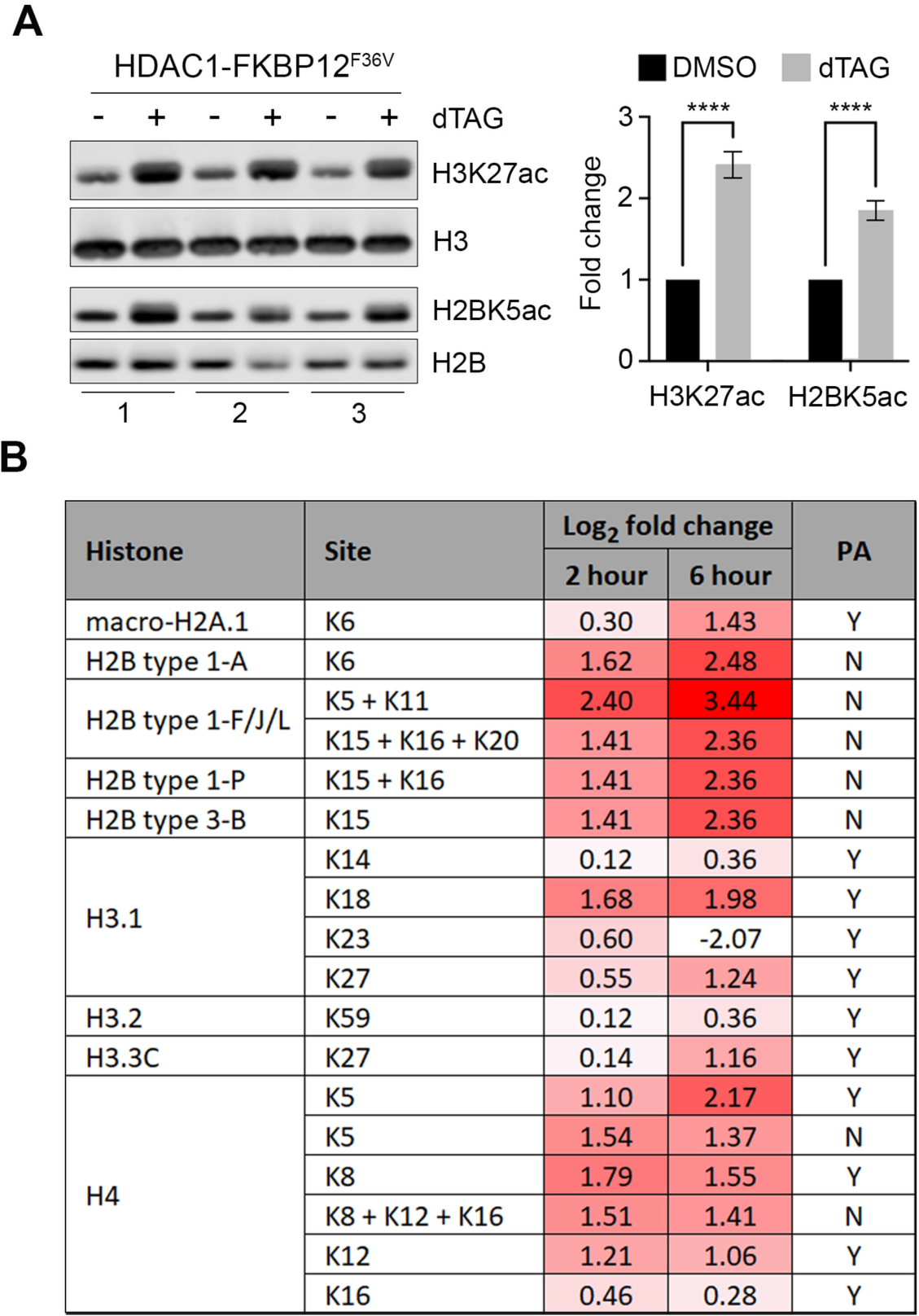
HDAC1-FKBP12^F36V^ degradation causes a global increase in histone tail acetylation. (**A**) Western blots showing H3K27ac/H2BK5ac levels -/+ 50 nM dTAG-13. Graph shows the relative changes normalized to total H3/H2B (n=3 +/-SD) (**** P<0.0001). (**B**) Table showing the log_2_ fold change in acetylation at the indicated histone sites following 2 or 6 hours of 50 nM dTAG-13 treatment detected via label-free quantitative mass spectrometry. Where multiple sites are indicated e.g. K5 + K11 the acetylation of multiple lysines was detected on the same peptide. PA column indicates whether samples were treated with propionic anhydride before trypsin digest to block unmodified lysine residues to generate peptides suitable for analysis.

### HDAC1 is critical for maintaining the gene expression network of ESCs

Due to the significant and rapid increases in histone acetylation, we reasoned that these would likely result in commensurate changes in gene expression. We performed RNA-seq at 2, 6 and 24 hours of dTAG treatment vs a DMSO control (with endogenous HDAC1/2 removed). Within 2 hours there were 275 upregulated and 15 downregulated genes (Fig 3A, full lists of gene expression changes in Supplementary table S3). 95% of the differentially expressed genes (DEGs) at 2 hours were upregulated, which fits with the canonical role of HDACs as transcriptional repressors. However, the absence of downregulated genes at this timepoint is also likely reflective of mRNA stability and the strict cut-offs that we applied (baseMean >50, padj <0.01, fold change >2). Over 1,500 DEGs (1,153 up vs 443 down) were identified at 6 hours, with the proportion of downregulated genes increasing to 28%. The increasing number of DEGs highlights that some genes respond more slowly to HDAC1 removal. Additionally, some changes at this point may be due to the downstream effects exerted by the genes that are upregulated at 2 hours. By 24 hours there are 1,446 upregulated genes with 967 downregulated (40% of total). The increasing proportion of downregulated genes suggests that the absence of HDAC1 results in the inability to maintain the necessary recruitment of RNA polymerase II (RNAP II) to these genes leading to the downregulation of their expression. Additionally, at 24 hours some of the gene expression changes that are seen will be associated with the cell death noted on Fig 1B and 1C.

**Figure 3.**
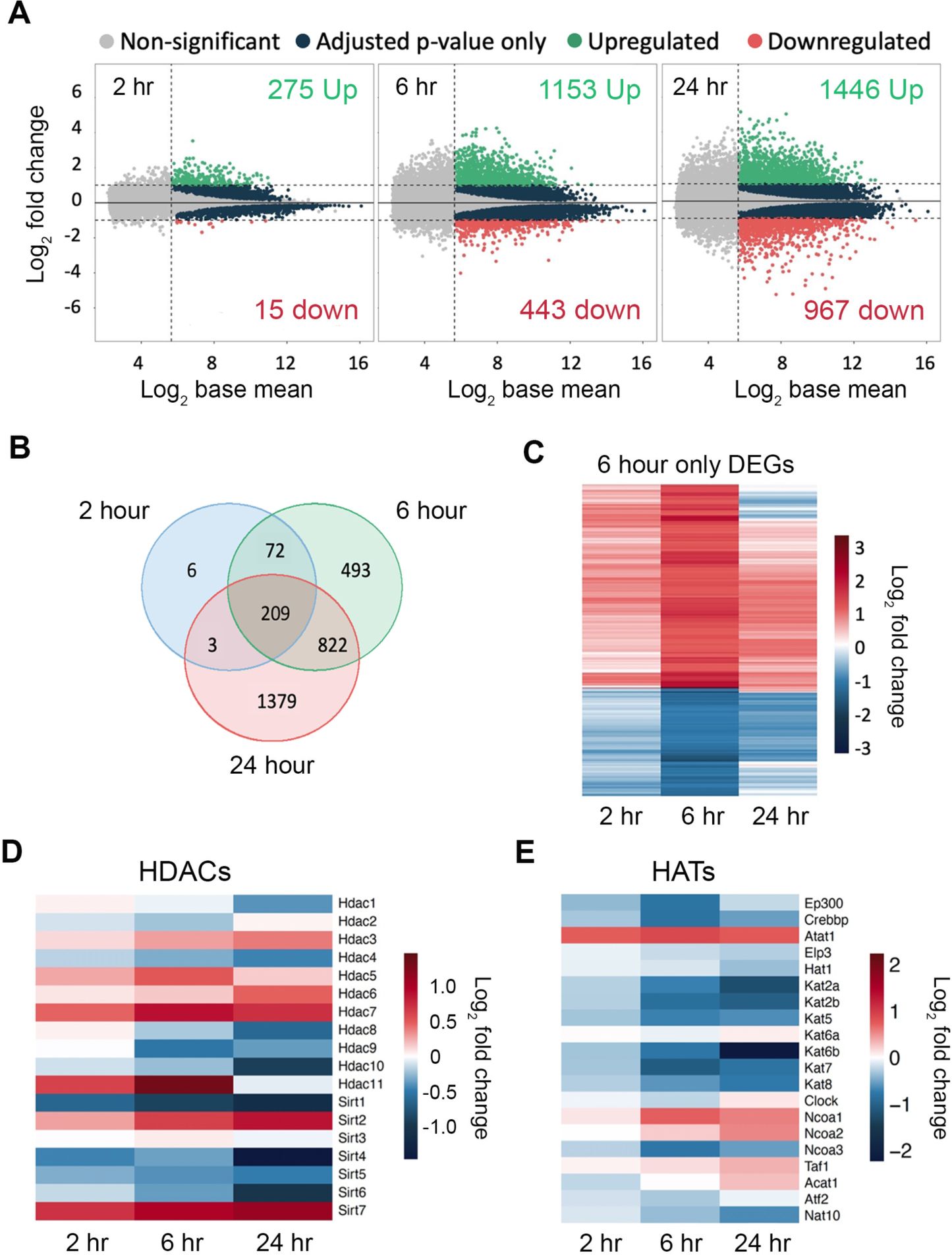
The ESC transcriptome is rapidly dysregulated following HDAC1-FKBP12^F36V^ degradation. (**A**) MA plots showing the numbers of differentially expressed genes (DEGs) following 2, 6 and 24 hours of treatment with 100nM dTAG^V^-1. DEGs defined as: p-adjusted value of <0.01, log_2_ fold change of > +/-1 and an average expression (baseMean) >50 normalised counts. (**B**) Venn diagram showing the overlaps of DEGs from (A) at the indicated time points. (**C**) Heatmap showing the log_2_ fold change of the 493 DEGs exclusive to the 6 hour time point on (B). (**D**) Heatmap showing the log_2_ fold change of the expression of Histone Deacetylases at the indicated time points. (**E**) Heatmap showing the log_2_ fold change of the expression of Histone Acetyltransferases at the indicated time points.

There is a strong overlap of the 290 DEGs at 2 hours with those at 6 and 24 (97% - Fig 3B) indicating that genes which are primary targets of HDAC1 remain affected at later time points and there is no compensatory mechanism to rescue their expression levels. In contrast, 30% of the DEGs (493) at 6 hours were only found at this time point (exemplar up and downregulated genes are shown in Supplementary Fig S4), where initial changes in expression are brought close to their original levels by 24 hours. The distribution of these genes (322 up and 171 down) was approximately the same as the overall distribution at 6 hours (Fig 3C). Possible explanations for these genes returning below the cut-offs for significance include compensation through increased activity of other HDACs or a change in the balance between HAT and HDAC activity. Fig 3D shows that there were some modest increases in the transcript levels of other nuclear HDACs. HDAC3 is the most obvious candidate to compensate for the loss of HDAC1/2 and there was a moderate increase in *Hdac3* RNA levels (Supplementary Fig S5A). However, there was no increase in HDAC3 protein levels at any time point (Supplementary Fig S5B and S5C). Interestingly, the expression levels of the Sirtuins (class III NAD^+^-dependent HDACs) appear more sensitive to the loss of HDAC1/2. *Sirt2* and particularly *Sirt7* showed increased expression. SIRT7 depletion does not result in a global change in the acetylation of the nucleolar or nuclear proteome suggesting that it is not responsible for returning the expression of these genes to normal levels (41-43). Among the HATs there is an overall pattern of downregulation (Fig 3E), including the key enzymes p300 and CBP (*Ep300* and *Crebbp* in Fig 3E). Therefore, it may be that reduced HAT activity results in the ‘6 hour only’ genes returning below the cut-offs for DEGs by 24 hours.

Gene Ontology (GO) analysis was performed on the genes upregulated at 2 hours to determine the biological processes that were directly affected by HDAC1 removal. Nervous system development is the GO term with the most associated upregulated genes (Fig 4A), suggesting that this network of genes is becoming derepressed in the absence of HDAC1. Several of the terms highlighted were in reference to upregulated genes linked to the circadian clock (Fig 4A). The circadian clock is controlled by an oscillating feedback loop involving the key factors *Bmal1* (*Arntl*), *Clock*, *Per* and *Cry* (44). Despite expressing the clock factors, ESCs do not have a functional clock system (45); it is thought that the clock factors may help to regulate proliferation in ESCs a role which they play in differentiated cells (reviewed in (44)). The lack of a functional clock system in ESCs is highlighted here as despite strong increases in *Per1*, *Per2* and *Cry2* at 6 hours (Fig 4B), there is not a reduction in *Arntl* or *Clock* as would be expected with a functional clock system (Fig 4B). As only 15 genes were downregulated at 2 hours, we performed GO analysis on the genes downregulated at 6 hours to determine the processes most directly affected. The most enriched terms at this time point included cellular response to LIF and positive regulation of stem cell population maintenance (Fig 4C). When we examined the broader network of pluripotency-associated factors, we noticed a general downward trend in their expression (Supplementary Fig S6). Rapid degradation of HDAC1 via the dTAG correlates well with previous studies (9, 46) but the downregulation of pluripotency-associated factors becomes apparent far more quickly than using conditional KOs. In particular, we noted downregulation of the pluripotency-associated factors *Prdm14*, *Tet1* and *Tet2* (Fig 4D). PRDM14, TET1 and TET2 are important in maintaining low DNA methylation levels in ESCs (47, 48). We observed a reciprocal change in genes associated with DNA methylation, including increased expression of the de novo DNA methyltransferases, *Dnmt3a* and *Dnmt3b* (Fig 4D). Rapid and specific degradation of HDAC1 using the dTAG system has allowed us to identify direct transcriptional targets of HDAC1 for the first time, revealing critical roles in the pluripotent gene network, and regulation of the DNA methylation machinery.

**Figure 4.**
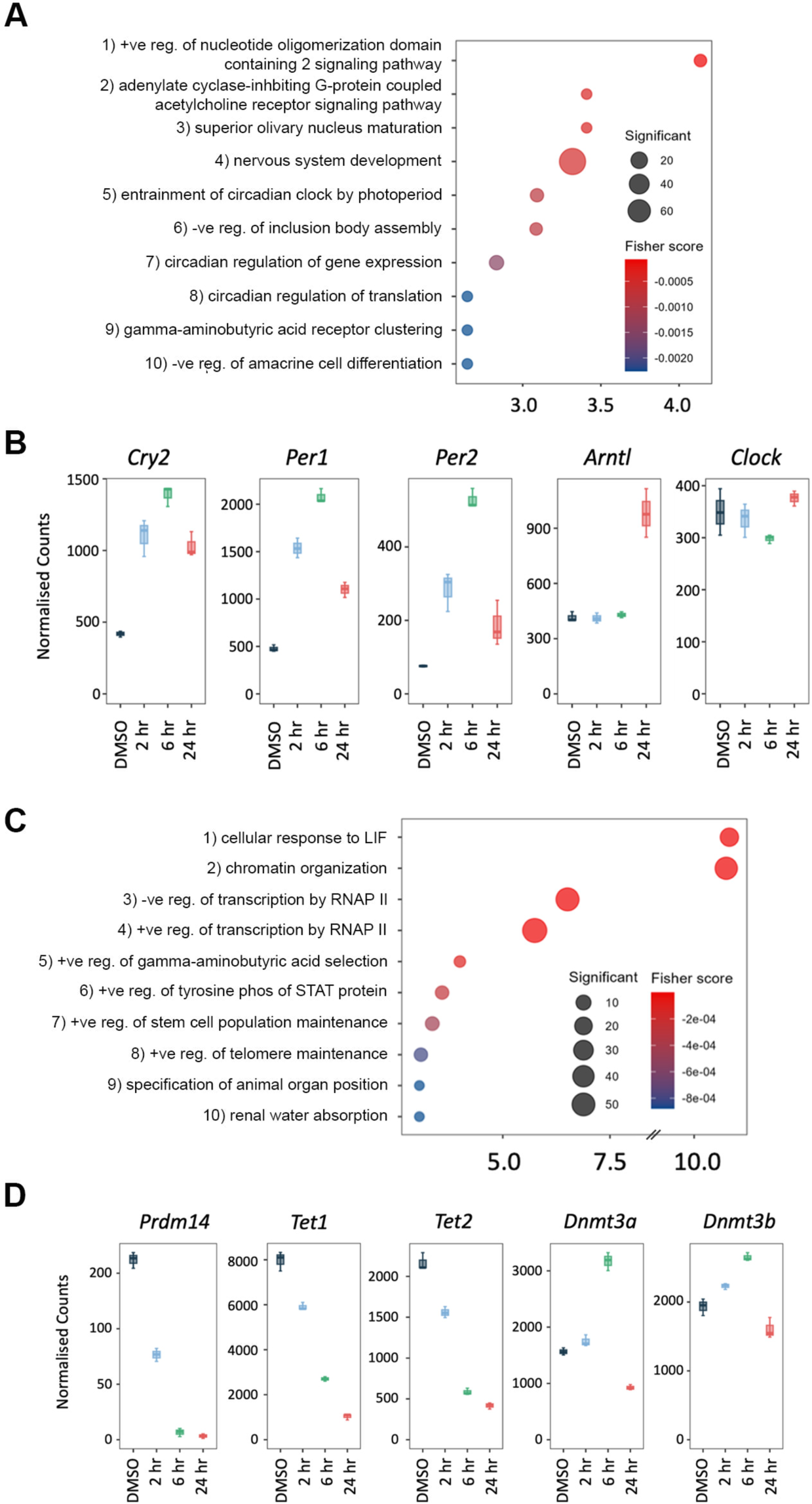
HDAC1-FKBP12^F36V^ degradation causes rapid changes in genes associated with ESC characteristics. (**A**) The 10 most enriched biological process gene ontology (GO) terms associated with the upregulated genes (padj <0.01, log2 fold change > +1) following 2 hours of HDAC1-FKBP12^F36V^ degradation. (**B**) Boxplots displaying the normalised count values of genes associated with circadian rhythm at the indicated time points. (**C**) The 10 most enriched biological process GO terms associated with the downregulated genes (padj <0.01, log2 fold change > -1) following 6 hours of HDAC1-FKBP12^F36V^ degradation. (**D**) Boxplots displaying the normalised count values of genes associated with DNA methylation at the indicated timepoints.

### HDAC1 degradation causes genome wide changes in histone acetylation that underpin changes in the transcriptome

To examine loci-specific changes in histone acetylation following HDAC1 loss we performed ChIP-seq for H2BK5ac and H3K27ac following 6 hours of dTAG treatment. We reasoned that these data should allow us to determine how alterations in histone acetylation affect the expression of HDAC1 sensitive genes identified by RNA-seq. H2BK5ac and H3K27ac were chosen because both mark active enhancers and promoters and were strongly increased following HDAC1 degradation at the same timepoint (Fig 2A and 2B). We first looked at upregulated genes identified by RNA-seq, for *Cep170b* (Fig 5A) and *Mast* 3 (Fig 5B) there was increased ChIP signal following dTAG treatment for both H2BK5ac and H3K27ac. There was a notable increase in H2BK5ac and H3K27ac around the transcription start site (TSS) (regions highlighted in blue on Fig 5A and 5B), including the promoters of these genes. Metaplots of all upregulated genes showed that H3K27ac and H2BK5ac were increased at the promoter and gene bodies following HDAC1 removal (Fig 5C). However, this trend is restricted to upregulated genes as we observed only modest increases (H3K27ac) or no change (H2BK5ac) in acetylation when we expanded the analysis to include all genes (Supplementary Fig S7A). Peak calling for H3K27ac revealed that 17,123 sites showed increased H3K27ac while 3,313 sites were reduced following dTAG treatment (FDR<=0.05, Fig 5D). The greatest proportion of upregulated H3K27ac peaks were indeed found in promoter regions (Fig 5E). More than 75% (881 of 1153) of upregulated genes identified by RNA-seq have at least one upregulated H3K27ac peak (Fig 5F). These results indicate that the increased acetylation in the proximity of upregulated genes (particularly at the promoter) is a general feature of increased gene expression within 6 hours of HDAC1 degradation. We were unable to perform a similar analysis with the H2BK5ac dataset as an overall broadening of the acetyl peak prevented accurate peak calling (Supplementary Fig S7B). However, we did also observe an increase in the average H2BK5ac and H3K27ac signal across promoters of upregulated genes (Fig 5G). Rapid degradation of HDAC1 is here shown to lead to an increase in acetylation specifically at genes which are upregulated in response to HDAC1 degradation.

**Figure 5.**
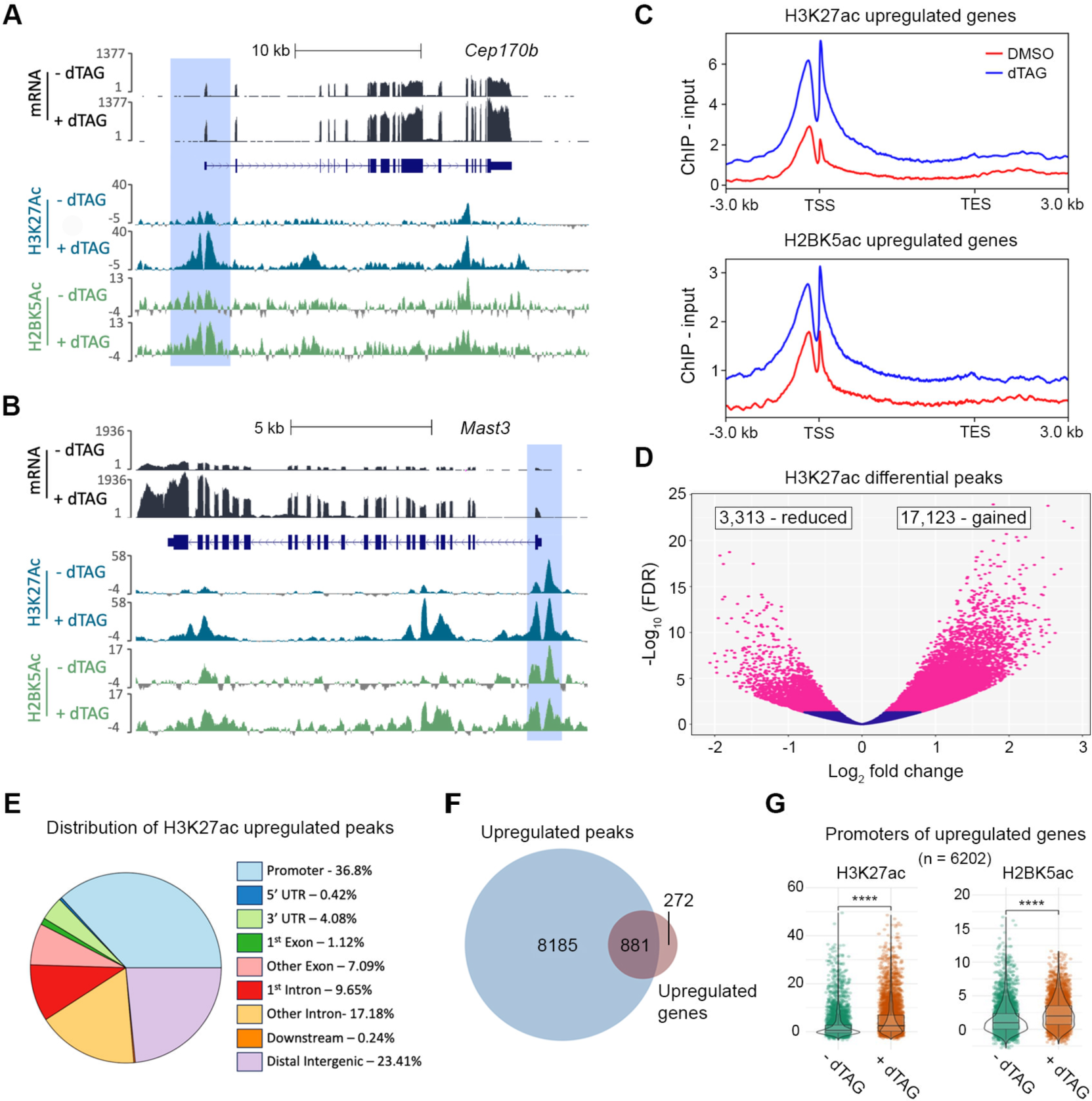
Upregulated genes show increased acetylation at the transcriptional start site following HDAC1-FKBP12^F36V^ degradation. (**A, B**) Tracks from the UCSC genome browser (38) showing the effect of 6 hours of 100 nM dTAG^V^-1 treatment on the mRNA, H3K27ac and H2BK5ac levels at the *Cep170b* (A) and *Mast3* (B) loci. Regions around the transcription start site (TSS) are highlighted. (**C**) Metaplots showing the average signal (ChIP - Input) for H3K27ac and H2BK5ac following 6 hours of 100 nM dTAG^V^-1 or DMSO treatment as indicated across the upregulated genes determined by RNA-seq (padj <0.01, log2 fold change > +1), including the regions -/+ 3 kb from the transcription start site (TSS) and transcription end site (TES). (**D**) Volcano plot showing the differential peaks for H3K27ac identified by ChIP-seq following 6 hours of 100 nM dTAG^V^-1 treatment. (**E**) Plot indicating the percentage of the upregulated peaks defined on (D) associated with the indicated genomic features. (**F**) Venn diagram indicating the overlap between the upregulated genes defined by RNA-seq and the upregulated H3K27ac peaks (where peaks are assigned to the closest gene by proximity) identified on (D) following 6 hours of dTAG^V^-1 treatment. (**G**) Violin plots showing the average signal (ChIP - Input) across the promoter region (defined as from the TSS to -500 bp) of the upregulated genes -/+ 6 hours of 100 nM dTAG^V^-1 treatment as indicated (**** P<0.0001).

Explaining why so many genes are downregulated following the removal of HDAC1 is more difficult given that the removal of acetyl marks is thought of as a repressive action. Inspection of the *Nanog* locus (Fig 6A) revealed that changes in acetylation around the TSS were less pronounced than those presented above for upregulated genes (Fig 5A and 5B). Metaplots for all downregulated genes showed that H3K27ac was unchanged around the TSS while there was only a small decrease in H2BK5ac levels (Fig 6B). These results suggest that a decrease in acetylation around the TSS is not responsible for downregulation of gene expression. Further inspection of the downregulated genes revealed that the 3 most strongly downregulated genes at 6 hours (*Prdm14*, *Inhbb* and *Nanog*) all have associated super-enhancers (SEs). SEs are regions of clustered enhancers that control expression of genes critical for cell identity in any given cell type (35). Perturbations of HDAC activity have previously been shown to have strong effects on SE dynamics (49, 50). Additional examination of the *Nanog* locus (Fig 6A) showed that there was a reduction in both H3K27ac and H2BK5ac at the associated SE. This is also true of the SEs associated with *Prdm14* and *Inhbb* (Supplementary Fig S8A and S8B). 11 of the top 50 most downregulated genes are regulated by SEs and this extends to ∼9% of all downregulated genes (Supplementary Fig S9A). In contrast, none of the top 50 most upregulated genes are SE regulated and less than 0.5% are in total (Supplementary Fig S9B). Metaplots across all SEs showed a clear downregulation of H3K27ac (Fig 6B), and an even more pronounced reduction in H2BK5ac that spreads beyond SE boundaries (Fig 6B).

**Figure 6.**
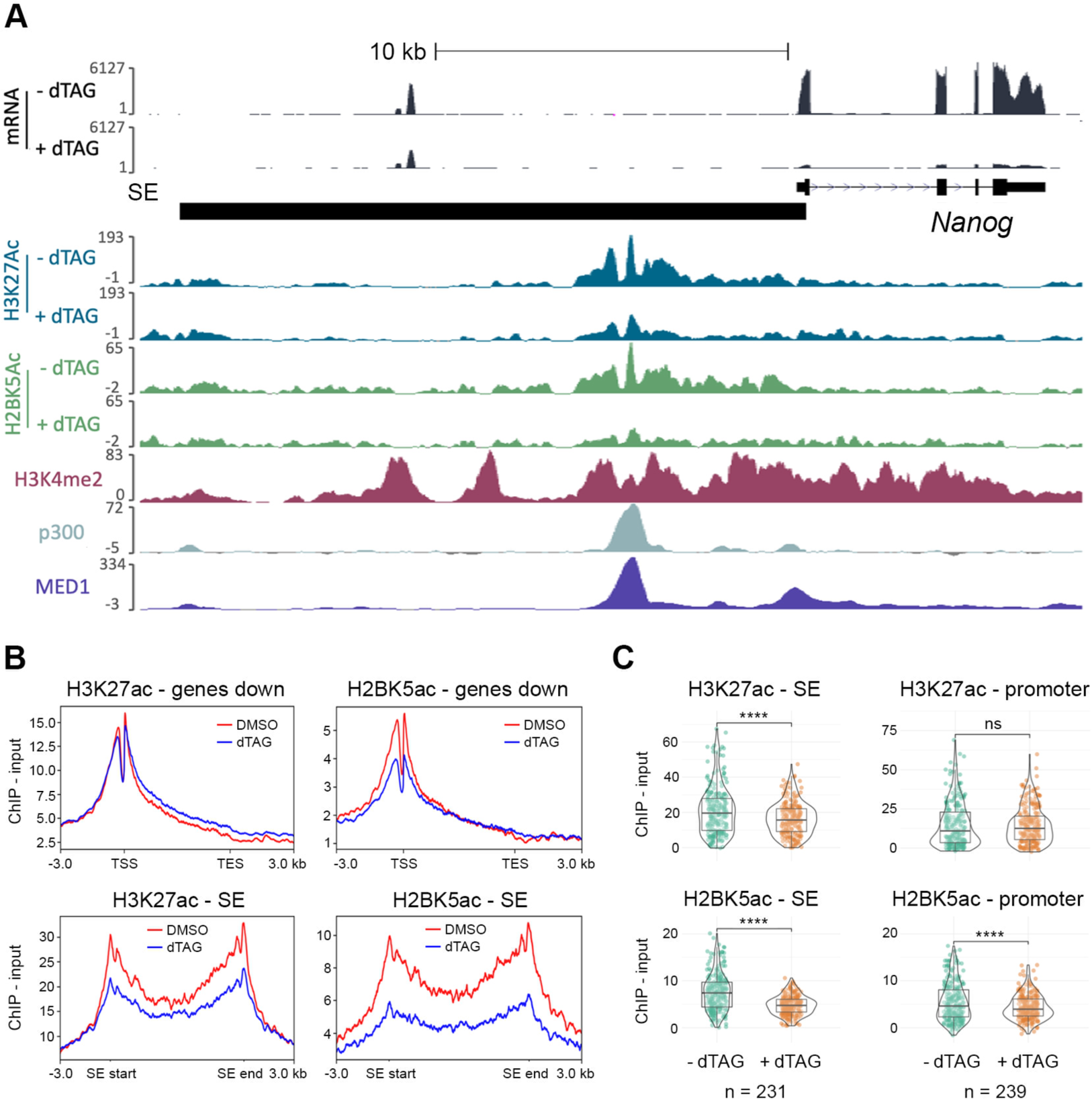
HDAC1-FKBP12^F36V^ degradation causes decreased acetylation at super-enhancers (SEs) and downregulation of SE-associated gene expression. (**A**) Tracks from the UCSC genome browser (38) showing the effect of 6 hours of 100 nM dTAG^V^-1 treatment on the mRNA, H3K27ac and H2BK5ac levels at the *Nanog* locus. SE region is indicated. H3K4me2 (produced in our lab previously) and p300/MED1 (36, 37) ChIP data are also shown, highlighting the SE region. (**B**) Metaplots showing the average signal (ChIP - Input) for H3K27ac and H2BK5ac following 6 hours of 100 nM dTAG^V^-1 or DMSO treatment as indicated across the downregulated genes determined by RNA-seq (padj <0.01, log2 fold change > -1) and SE regions, including the regions -/+ 3 kb from the transcription start site (TSS) and transcription end site (TES) or SE start/end as indicated. (**C**) Violin plots showing the average signal (ChIP - Input) -/+ 100 nM dTAG^V^-1 treatment as indicated across SE regions (defined in (35)) and the promoter regions of SE-associated genes (defined as from the TSS to -500 bp). Genes with overlapping SE and promoter regions were excluded from this analysis (ns = non-significant, **** P<0.0001).

To get a general view of the importance of HDAC1 activity at SE-dependent genes, we ranked the genes based on their overall change in expression and then compared the relative change in acetylation (H2BK5ac and H3K27ac) at the SE versus promoter (Table 1, see Supplementary Table S4 for full list). Manual curation of the 231 SE regions in ESCs (35) suggests that they regulate 206 genes (with some genes having more than one SE e.g., *Inhbb*). The top 20 most downregulated genes are presented in Table 1. Significantly, we observed a stronger reduction in H3K27ac at the SE for these genes than the promoter. For example, *Otx2* showed an approximately 8-fold reduction in expression with around a 27% decrease in H3K27ac at the SE but an increase in acetylation at the promoter (up 29%). *Tet2* showed a similar effect with an approximately 3.7-fold reduction in expression, H3K27ac reduced by 57% at the SE, but increased by 5% at the promoter. The same trend is also true of H2BK5ac, although for this modification we also observed a smaller but consistent decrease at the promoter (Table 1, see *Manba*, 59% decrease at the SE and 18% reduction at the promoter). Plotting relative H3K27ac and H2BK5ac levels across all SE-dependent genes (Fig 6C) revealed a difference between these two sites of histone acetylation. The H3K27ac signal was reduced at the majority of SEs, while promoters for the same genes showed no significant change. H2BK5ac on the other hand, was significantly reduced at both the SE and the promoter, although the relative decrease was greater at the SE. These results show the requirement for HDAC1 activity at SEs and highlight that a proportion of the genes most strongly downregulated by HDAC1 removal can be explained by an unexpected reduction in histone acetylation at the SE, with or without a change in acetylation at the promoter.

**Table 1.** Table showing the change in acetylation at the super-enhancers (SEs) and promoters of the 20 most strongly downregulated SE-associated genes. Columns from left to right: ranking of genes by expression change (from most to least downregulated), the gene to which we have assigned the SE here based on manual interpretation of RNA-seq and ChIP-seq data (pluripotency-associated genes in the top 10 highlighted), the log_2_ fold change value for expression by RNA-seq following 6 hours of HDAC1-FKBP12^F36V^ degradation, the gene the SE was originally assigned to (defined in (35)), whether we have reassigned the SE to a new gene (reassigned SEs highlighted), the chromosome on which each SE is located followed by the start and end site, the percentage change of H3K27ac and H2BK5ac across the SE and promoter regions (of the genes that we have assigned). The expression and % change columns are colored with blue indicating a decrease and red indicating an increase, SEs/promoters that showed a > 200% change were omitted and are shown in grey.

| Ranking | New SE designation | Expression log<sub>2</sub> FC | Original SE designation | Reassigned | Chromosome | Start | End | H3K27ac SE (% change) | H2BK5ac SE (% change) | H3K27ac Prom (% change) | H2BK5ac Prom (% change) |
| --- | --- | --- | --- | --- | --- | --- | --- | --- | --- | --- | --- |
| 1 | Prdm14 | -4.02 | Prdm14 | No | chr1 | 13059534 | 13104684 | -40.90 | -33.92 | -91.48 | -32.94 |
| 2 | Inhbb (a) | -3.34 | Inhbb (a) | No | chr1 | 119304847 | 119305904 | 1.57 | -27.25 | 37.03 | 3.26 |
|  | Inhbb (b) | -3.34 | Inhbb (b) | No | chr1 | 119398508 | 119399454 | -63.13 | -44.10 | 37.03 | 3.26 |
| 3 | Nanog | -3.28 | Nanog | No | chr6 | 122690100 | 122707853 | -38.54 | -47.90 | -44.03 | -35.95 |
| 4 | Otx2 | -3.03 | Otx2 | No | chr14 | 48653438 | 48663525 | -26.94 | -12.89 | 29.36 | 127.33 |
| 5 | Gdf3 | -2.36 | Dppa3 | Yes | chr6 | 122662496 | 122664242 | -35.16 | -29.20 | 7.43 | -15.11 |
| 6 | H2-M5 | -2.30 | H2-M5 | No | chr17 | 36973257 | 36998051 | -10.98 | -36.68 | -18.63 | -58.58 |
| 7 | Zfp42 | -2.14 | Zfp42 | No | chr8 | 43320382 | 43321401 | -36.16 | -44.46 | -32.11 | -56.46 |
| 8 | Dmtn | -2.11 | Dmtn | No | chr14 | 70622852 | 70636123 | 18.84 | 14.17 | 30.55 | 46.49 |
| 9 | Tet2 | -1.87 | Tet2 | No | chr3 | 133518467 | 133534684 | -57.49 | -68.89 | 5.34 | -56.72 |
| 10 | Gbx2 | -1.85 | Gbx2 | No | chr1 | 89870372 | 89876952 | -38.18 | -54.61 | 1806.17 | 158.81 |
| 11 | Mir290-295 | -1.82 | Mir290a | No | chr7 | 3193003 | 3218182 | -31.60 | -46.28 | N/A | N/A |
| 12 | Mllt6 | -1.69 | Mllt6 | No | chr11 | 97656359 | 97662845 | 6.59 | -14.28 | -16.39 | -46.89 |
| 13 | Tbx3 (a) | -1.68 | Tbx3 (a) | No | chr5 | 119579640 | 119587054 | -40.11 | -47.17 | 13.94 | 10.36 |
|  | Tbx3 (b) | -1.68 | Tbx3 (b) | No | chr5 | 119679583 | 119721473 | -41.44 | -39.46 | 13.94 | 10.36 |
| 14 | Manba | -1.67 | Manba | No | chr3 | 135545992 | 135547780 | -45.52 | -58.68 | 179.89 | -18.10 |
| 15 | Tet1 | -1.55 | Tet1 | No | chr10 | 62883646 | 62898815 | 2.26 | -22.31 | 28.67 | -17.50 |
| 16 | Fgf4 | -1.51 | Fgf4 | No | chr7 | 144850967 | 144864811 | 11.73 | -23.26 | -9.93 | -25.78 |
| 17 | Zfp710 | -1.49 | Zfp710 | No | chr7 | 80015022 | 80025077 | 4.05 | -26.17 | 26.40 | -10.59 |
| 18 | Setd1b | -1.45 | Rhof | Yes | chr5 | 123134650 | 123140719 | -15.51 | -30.48 | 11.95 | -32.43 |
| 19 | Bvht | -1.44 | Mir145a | Yes | chr18 | 61627890 | 61628746 | -36.49 | -53.78 | -28.80 | -35.29 |
| 20 | Rbpj | -1.44 | Rbpj | No | chr5 | 53541938 | 53556088 | -2.80 | -32.93 | -19.87 | -32.57 |

## Discussion

HDAC1/2 are the dominant deacetylases in ESCs, contributing more than half of the total deacetylase activity (9). Furthermore, HDAC1 appears to be the dominant paralogue as the three predominant corepressor complexes (SIN3A, NuRD and CoREST) show reduced deacetylase activity in the absence of HDAC1 but not HDAC2 (26). It has been known for more than 30 years that inhibiting HDAC activity in cells produced a combination of up- and downregulated genes (51, 52). However, many of these initial experiments required treating cells with inhibitors for days at a time, or genetic KOs where the slow turnover of the enzymes and their associated complexes led to indirect effects that muddied the mechanistic waters. With the dTAG system allowing degradation of HDAC1-FKBP12^F36V^ within an hour (Fig 1A) we have been able to investigate HDAC1 activity on a timescale that has not previously been possible. HDAC1-FKBP12^F36V^ expression rescued the viability of *Hdac1/2* DKO ESCs (Fig 1B) highlighting that HDAC2 is not required and confirming previous findings that either paralog is sufficient in ESCs (26). However, the loss of viability following dTAG treatment in HDAC1-FKBP cells occurs 3 days earlier than using conditional knockouts (9), highlighting the power of the dTAG system.

The prevailing assumption regarding class I HDACs, is that they are recruited into numerous corepressor complexes (7 in total) to confer substrate specificity and guide the deacetylase activity to specific loci. As 6 of these 7 corepressor complexes contain HDAC1 (and/or HDAC2) it is no surprise that degradation of HDAC1 revealed a wide range of substrates across the core histones (Fig 2B). Many of the sites identified are high stoichiometry acetylation sites, including H4K5, H4K8, H4K12, H4K16, H2BK5 and H2BK20 (1, 53-56). Strikingly, several of the sites (including those on H2B) are targets for the crucial acetyltransferases, p300/CBP (39). The importance of these H2B sites has recently been realised, as indicators of enhancer strength and their intensity at promoters is predictive of CBP/p300 regulated genes (36). Acetylation sites on the H2B tail are normally subject to a very fast acetylation/deacetylation cycle, highlighted by their short half-lives (39). In the absence of HDAC1, p300/CBP have unrestricted access to acetylate these sites resulting in rapid increases (Fig 2B). The increases in H4K5ac and H4K12ac suggest that HDAC1 balances the activity of HAT1 which deposits these marks on newly synthesised histones (57). These results show that HDAC1 balances the acetyltransferase activity of the most active HATs, helping explain why HDAC1/2 are the most abundant and active HDACs in cells (9, 58).

RNA-seq analysis showed that the immediate consequence of HDAC1 degradation was rapid upregulation of gene expression with 275 out of 290 DEGs upregulated at 2 hours (Fig 3A). Given the timescale of derepression we would expect the majority of these genes to be *direct* targets of HDAC1-containing complexes. Our results mirror precision run-on sequencing (PRO-seq) results in drosophila, where 10 minutes of treatment with the HDAC inhibitor (HDACi) TSA caused 96 upregulated genes, but no genes were downregulated (59). PRO-seq allows a far faster readout but the trends observed do match. GO analysis revealed that nervous system development was one of the biological processes most strongly associated with the 2-hour upregulated genes (Fig 4A). This upregulation of genes associated with nervous system development may be due to the loss of repression normally mediated by the HDAC1/2 containing CoREST complex (60).

ChIP-seq revealed a particular increase in H3K27ac and H2BK5ac around the TSS of upregulated genes (Fig 5C). There is continued debate as to whether increased acetylation is a cause or consequence of increased transcription, with recent studies coming down on either side of the argument, promoting transcription via release of paused RNAP II (59), or on the contrary, that global acetylation is entirely dependent on transcription (61). Fig 5C reveals that both H2BK5ac and H3K27ac were increased immediately downstream of the TSS following dTAG treatment. These results are compatible with findings from Vaid *et al*, whereby they identified increased acetylation of the first nucleosomes past the TSS following TSA treatment, facilitating increased release of paused polymerase giving upregulation of these genes (59).

Despite the predominant effect on gene expression at 2 hours being upregulation, 15 genes were downregulated including the pluripotency-associated factor *Prdm14* (Fig 4D). PRDM14 maintains the low DNA methylation levels of ESCs by repressing expression of the de novo DNA methyltransferases, *Dnmt3a/b* (48) and by recruiting TET proteins to loci for DNA demethylation (47). At 6 hours the effects of reduced *Prdm14* expression were apparent as there was a greater than 2-fold increase in *Dnmt3a* expression with a more modest increase in *Dnmt3b* (Fig 4D). Additionally, *Tet1* and *Tet2* were downregulated more than 2-fold by 6 hours (Fig 4D). These results suggest that the typically low DNA methylation levels in ESCs would be perturbed. By 6 hours, 443 genes were downregulated, indicating that HDAC1 is also required for maintaining the expression of many genes (Fig 3A). GO analysis revealed that many of these genes were associated with stem cell maintenance (Fig 4C). This is consistent with our recent PRO-seq study in ESCs, using the pan HDAC-inhibitor, LBH589 (46). A 2 hour treatment with LBH589 caused down-regulation of the pluripotent gene network due to reduced RNAP II recruitment at promoters and super-enhancers (SEs). The data in this study strongly indicates that HDAC1 and its associated complexes are required for this activity.

The auto-regulatory gene networks that control cell identity (e.g., pluripotency in ESCs) are regulated by SEs (35). In the last few years, a number of studies have demonstrated that SEs are dependent on HDAC activity, using a range of HDAC inhibitors, e.g., largazole and MS-275 (49, 50, 62). In this study, we have manually curated the 231 known SE regions in ESCs, which regulate 206 genes (some genes have more than one SE, e.g., *Mycn* has 3)(Supplementary Table 4). 98 of 206 genes are reduced >1.5-fold following loss of HDAC1, with only 9 increased, a ratio of 10:1 in favour of reduced activity. The 98 downregulated genes collectively have 113 SEs, with 93% (105/113) showing reduced H2BK5ac signal. 7 of the top-10 most strongly downregulated SE-associated genes are pluripotency-associated factors (Table 1 genes highlighted in green: *Prdm14, Nanog, Otx2, Gdf3, Zfp42, Tet2 and Gbx2*). *Nanog* and *Prdm14* were downregulated more than 9-fold at 6 hours and had reduced acetylation at their SEs (Fig 6A and Supplementary Fig S8A). Downregulation of the extended pluripotency network at 24 hours (Supplementary Fig S6 – e.g., *Utf1*, *Lefty1/2*, *Nodal*, *Dppa4*, *Tcl1*) may occur because of reduced NANOG and ESRRB binding at regulatory elements. Given the correlation between HDAC1-sensitivity and reduced H3K27ac/H2BK5ac signal, we were able to reassign the gene targets for 51 SEs (22%) to hopefully produce a definitive list of SE-dependent genes in ESCs (Supplementary Table S4).

In conclusion, our results have shown that HDAC1 regulates the majority of acetylation sites on the core histones, with lysine residues on H2B proving the most sensitive to loss of HDAC1. The greatly increased temporal resolution of the dTAG system allowed us to study the early consequences of HDAC1 degradation on gene expression. As early as 2 hours we monitored derepression of the gene network regulating circadian rhythm. While most of the early genes were upregulated, many of the most HDAC1-sensitive genes were downregulated, many of these genes form part of the pluripotency-associated gene network. The core regulatory network in ESCs is paradoxically reliant on HDAC1 to maintain histone acetylation levels at SEs, presumably in a dynamic relationship with histone acetyltransferases, such as p300. An acetylation/deacetylation cycle is likely a key part of active transcription (63) with inhibition of either activity a *stick in the spokes* for the overall activity of RNAP II.

## Data Availability

All sequencing data in this study has been deposited to GEO under the SuperSeries GSE269448. The mass spectrometry proteomics data have been deposited to the ProteomeXchange Consortium via the PRIDE (32) partner repository with the dataset identifier PXD053032. Cell lines and plasmids used in this study are available upon request.

## Acknowledgements

Thank you to members of the Cowley and Schwabe groups for critical comments and feedback on the data throughout the project. We thank Dr. Nicolas Sylvius and the NUCLEUS genomics facility for help with RNA extractions and quality checking of RNA/DNA. Plasmid constructs were generated by the PROTEX facility at the University of Leicester. D.M.E and S.N.L were supported by funding from the Medical Research Council (MRC) (MR/W00190X/1). K.A.S is part of the MRC IMPACT (Integrated Midlands Partnership for Biomedical Training) programme. I.M.B was supported by a Biotechnology and Biological Sciences Research Council (BBSRC) studentship from the Midlands Integrative Biosciences Training Partnership (MIBTP). S.M.C was supported by project grants from the MRC (MR/W00190X/1) and BBSRC (BB/P021689/1). MOC was supported by project grants from the MRC (MR/X012220/1, MR/W00190X/1).

## Author Contributions

D.M.E and K.A.S designed and performed wet-lab experiments and bioinformatics analysis. S.N.L conducted bioinformatics analysis. T.K.P and M.O.C performed mass-spectrometry experiments and analysis of histone acetylation. S.M.C designed the study and wrote the manuscript with input from all authors.

## References

1. Hansen BK, Gupta R, Baldus L, Lyon D, Narita T, Lammers M, et al. Analysis of human acetylation stoichiometry defines mechanistic constraints on protein regulation. Nat Commun. 2019 -3-5;10.

2. Barnes CE, English DM, Cowley SM. Acetylation & Co: an expanding repertoire of histone acylations regulates chromatin and transcription. Essays Biochem. 2019 04 23,;63(1):97-107.

3. Bannister AJ, Kouzarides T. Regulation of chromatin by histone modifications. Cell Res. 2011 -03;21(3):381-95.

4. Zaware N, Zhou M. Bromodomain biology and drug discovery. Nat Struct Mol Biol. 2019 - 10;26(10):870-9.

5. Zhang R, Erler J, Langowski J. Histone Acetylation Regulates Chromatin Accessibility: Role of H4K16 in Inter-nucleosome Interaction. Biophys J. 2017 -02-07;112(3):450-9.

6. Drazic A, Myklebust LM, Ree R, Arnesen T. The world of protein acetylation. Biochim Biophys Acta. 2016 -10;1864(10):1372-401.

7. Newman JC, He W, Verdin E. Mitochondrial protein acylation and intermediary metabolism: regulation by sirtuins and implications for metabolic disease. J Biol Chem. 2012 -12-14;287(51):42436-43.

8. Zupkovitz G, Tischler J, Posch M, Sadzak I, Ramsauer K, Egger G, et al. Negative and positive regulation of gene expression by mouse histone deacetylase 1. Mol Cell Biol. 2006 - 11;26(21):7913-28.

9. Jamaladdin S, Kelly RDW, O’Regan L, Dovey OM, Hodson GE, Millard CJ, et al. Histone deacetylase (HDAC) 1 and 2 are essential for accurate cell division and the pluripotency of embryonic stem cells. Proc Natl Acad Sci U S A. 2014 Jul 08,;111(27):9840-5.

10. Baker IM, Smalley JP, Sabat KA, Hodgkinson JT, Cowley SM. Comprehensive Transcriptomic Analysis of Novel Class I HDAC Proteolysis Targeting Chimeras (PROTACs). Biochemistry. 2023 Feb 7;62(3):645–56.

11. Morii A, Inazu T. HDAC8 is implicated in embryoid body formation via canonical Hedgehog signaling and regulates neuronal differentiation. Biochem Biophys Res Commun. 2022 -11-12;629:78-85.

12. Hassig CA, Fleischer TC, Billin AN, Schreiber SL, Ayer DE. Histone deacetylase activity is required for full transcriptional repression by mSin3A. Cell. 1997 May 02,;89(3):341-7.

13. Laherty CD, Yang WM, Sun JM, Davie JR, Seto E, Eisenman RN. Histone deacetylases associated with the mSin3 corepressor mediate mad transcriptional repression. Cell. 1997 May 02,;89(3):349-56.

14. Xue Y, Wong J, Moreno GT, Young MK, Côté J, Wang W. NURD, a novel complex with both ATP-dependent chromatin-remodeling and histone deacetylase activities. Mol Cell. 1998 Dec;2(6):851–61.

15. You A, Tong JK, Grozinger CM, Schreiber SL. CoREST is an integral component of the CoREST-human histone deacetylase complex. Proc Natl Acad Sci U S A. 2001 -2-13;98(4):1454-8.

16. Bantscheff M, Hopf C, Savitski MM, Dittmann A, Grandi P, Michon A, et al. Chemoproteomics profiling of HDAC inhibitors reveals selective targeting of HDAC complexes. Nat Biotechnol. 2011 Mar;29(3):255–65.

17. Itoh T, Fairall L, Muskett FW, Milano CP, Watson PJ, Arnaudo N, et al. Structural and functional characterization of a cell cycle associated HDAC1/2 complex reveals the structural basis for complex assembly and nucleosome targeting. Nucleic Acids Res. 2015 Feb 27,;43(4):2033-44.

18. Ding Z, Gillespie LL, Paterno GD. Human MI-ER1 alpha and beta function as transcriptional repressors by recruitment of histone deacetylase 1 to their conserved ELM2 domain. Mol Cell Biol. 2003 Jan;23(1):250–8.

19. Plaster N, Sonntag C, Schilling TF, Hammerschmidt M. REREa/Atrophin-2 interacts with histone deacetylase and Fgf8 signaling to regulate multiple processes of zebrafish development. Dev Dyn. 2007 Jul;236(7):1891–904.

20. Kelly RDW, Cowley SM. The physiological roles of histone deacetylase (HDAC) 1 and 2: complex co-stars with multiple leading parts. Biochem Soc Trans. 2013 Jun;41(3):741–9.

21. Dovey OM, Foster CT, Conte N, Edwards SA, Edwards JM, Singh R, et al. Histone deacetylase 1 and 2 are essential for normal T-cell development and genomic stability in mice. Blood. 2013 Feb 21,;121(8):1335-44.

22. Wang Z, Zang C, Cui K, Schones DE, Barski A, Peng W, et al. Genome-wide mapping of HATs and HDACs reveals distinct functions in active and inactive genes. Cell. 2009 Sep 4;138(5):1019–31.

23. Saha N, Liu M, Gajan A, Pile LA. Genome-wide studies reveal novel and distinct biological pathways regulated by SIN3 isoforms. BMC Genomics. 2016 Feb 13;17:111–5.

24. Choudhary C, Kumar C, Gnad F, Nielsen ML, Rehman M, Walther TC, et al. Lysine acetylation targets protein complexes and co-regulates major cellular functions. Science. 2009 Aug 14,;325(5942):834-40.

25. Schölz C, Weinert BT, Wagner SA, Beli P, Miyake Y, Qi J, et al. Acetylation site specificities of lysine deacetylase inhibitors in human cells. Nature Biotechnology. 2015 -04;33(4):415-23.

26. Dovey OM, Foster CT, Cowley SM. Histone deacetylase 1 (HDAC1), but not HDAC2, controls embryonic stem cell differentiation. Proceedings of the National Academy of Sciences of the United States of America. 2010 4 May;107(18):8242.

27. Nabet B, Roberts JM, Buckley DL, Paulk J, Dastjerdi S, Yang A, et al. The dTAG system for immediate and target-specific protein degradation. Nature Chemical Biology. 2018 - 05;14(5):431-41.

28. Nabet B, Ferguson FM, Seong BKA, Kuljanin M, Leggett AL, Mohardt ML, et al. Rapid and direct control of target protein levels with VHL-recruiting dTAG molecules. Nat Commun. 2020 Sep 18;11(1):4687-w.

29. Barnes C, English D, Broderick M, Collins M, Cowley SM. Proximity-dependent biotin identification (BioID) reveals a dynamic LSD1-CoREST interactome during embryonic stem cell differentiation. Mol Omics. 2021 -10-14.

30. Sidoli S, Kori Y, Lopes M, Yuan Z, Kim HJ, Kulej K, et al. One minute analysis of 200 histone posttranslational modifications by direct injection mass spectrometry. Genome Res. 2019 -6;29(6):978-87.

31. Cox J, Mann M. MaxQuant enables high peptide identification rates, individualized p.p.b.-range mass accuracies and proteome-wide protein quantification. Nat Biotechnol. 2008 -12;26(12):1367-72.

32. Perez-Riverol Y, Bai J, Bandla C, García-Seisdedos D, Hewapathirana S, Kamatchinathan S, et al. The PRIDE database resources in 2022: a hub for mass spectrometry-based proteomics evidences. Nucleic Acids Res. 2022 -01-07;50(D1):D543-52.

33. Fursova NA, Blackledge NP, Nakayama M, Ito S, Koseki Y, Farcas AM, et al. Synergy between Variant PRC1 Complexes Defines Polycomb-Mediated Gene Repression. Mol Cell. 2019 -06-06;74(5):1020,1036.e8.

34. Karolchik D, Hinrichs AS, Furey TS, Roskin KM, Sugnet CW, Haussler D, et al. The UCSC Table Browser data retrieval tool. Nucleic Acids Res. 2004 -01-01;32(Database issue):493.

35. Whyte WA, Orlando DA, Hnisz D, Abraham BJ, Lin CY, Kagey MH, et al. Master Transcription Factors and Mediator Establish Super-Enhancers at Key Cell Identity Genes. Cell. 2013 -4-11;153(2):307-19.

36. Narita T, Higashijima Y, Kilic S, Liebner T, Walter J, Choudhary C. Acetylation of histone H2B marks active enhancers and predicts CBP/p300 target genes. Nat Genet. 2023 Apr;55(4):679–92.

37. Narita T, Ito S, Higashijima Y, Chu WK, Neumann K, Walter J, et al. Enhancers are activated by p300/CBP activity-dependent PIC assembly, RNAPII recruitment, and pause release. Mol Cell. 2021 -05-20;81(10):2166,2182.e6.

38. Kent WJ, Sugnet CW, Furey TS, Roskin KM, Pringle TH, Zahler AM, et al. The human genome browser at UCSC. Genome Res. 2002 -06;12(6):996-1006.

39. Weinert BT, Narita T, Satpathy S, Srinivasan B, Hansen BK, Schölz C, et al. Time-Resolved Analysis Reveals Rapid Dynamics and Broad Scope of the CBP/p300 Acetylome. Cell. 2018 Jun 28;174(1):231,244.e12.

40. Creyghton MP, Cheng AW, Welstead GG, Kooistra T, Carey BW, Steine EJ, et al. Histone H3K27ac separates active from poised enhancers and predicts developmental state. Proc Natl Acad Sci U S A. 2010 -12-14;107(50):21931-6.

41. Michishita E, Park JY, Burneskis JM, Barrett JC, Horikawa I. Evolutionarily conserved and nonconserved cellular localizations and functions of human SIRT proteins. Mol Biol Cell. 2005 Oct;16(10):4623–35.

42. Tsai Y, Greco TM, Cristea IM. Sirtuin 7 plays a role in ribosome biogenesis and protein synthesis. Mol Cell Proteomics. 2014 Jan;13(1):73–83.

43. Kiran S, Anwar T, Kiran M, Ramakrishna G. Sirtuin 7 in cell proliferation, stress and disease: Rise of the Seventh Sirtuin! Cell Signal. 2015 Mar;27(3):673-82.

44. Dierickx P, Van Laake LW, Geijsen N. Circadian clocks: from stem cells to tissue homeostasis and regeneration. EMBO Rep. 2018 -01;19(1):18-28.

45. Yagita K, Horie K, Koinuma S, Nakamura W, Yamanaka I, Urasaki A, et al. Development of the circadian oscillator during differentiation of mouse embryonic stem cells in vitro. Proc Natl Acad Sci U S A. 2010 -02-23;107(8):3846-51.

46. Kelly RDW, Stengel KR, Chandru A, Johnson LC, Hiebert SW, Cowley SM. Histone deacetylases maintain expression of the pluripotent gene network via recruitment of RNA polymerase II to coding and noncoding loci. Genome Res. 2024 Feb 7;34(1):34–46.

47. Okashita N, Kumaki Y, Ebi K, Nishi M, Okamoto Y, Nakayama M, et al. PRDM14 promotes active DNA demethylation through the ten-eleven translocation (TET)-mediated base excision repair pathway in embryonic stem cells. Development. 2014 -01;141(2):269-80.

48. Leitch HG, McEwen KR, Turp A, Encheva V, Carroll T, Grabole N, et al. Naive pluripotency is associated with global DNA hypomethylation. Nat Struct Mol Biol. 2013 -03;20(3):311-6.

49. Sanchez GJ, Richmond PA, Bunker EN, Karman SS, Azofeifa J, Garnett AT, et al. Genome-wide dose-dependent inhibition of histone deacetylases studies reveal their roles in enhancer remodeling and suppression of oncogenic super-enhancers. Nucleic Acids Res. 2018 -2-28;46(4):1756-76.

50. Gryder BE, Wu L, Woldemichael GM, Pomella S, Quinn TR, Park PMC, et al. Chemical genomics reveals histone deacetylases are required for core regulatory transcription. Nat Commun. 2019 -07-08;10(1):1-12.

51. Clayton AL, Hazzalin CA, Mahadevan LC. Enhanced histone acetylation and transcription: a dynamic perspective. Mol Cell. 2006 -08-04;23(3):289-96.

52. Smith CL. A shifting paradigm: histone deacetylases and transcriptional activation. Bioessays. 2008 -01;30(1):15-24.

53. Abshiru N, Caron-Lizotte O, Rajan RE, Jamai A, Pomies C, Verreault A, et al. Discovery of protein acetylation patterns by deconvolution of peptide isomer mass spectra. Nat Commun. 2015 -10-15;6:8648.

54. Feller C, Forné I, Imhof A, Becker PB. Global and specific responses of the histone acetylome to systematic perturbation. Mol Cell. 2015 -02-05;57(3):559-71.

55. Zhou T, Chung Y, Chen J, Chen Y. Site-Specific Identification of Lysine Acetylation Stoichiometries in Mammalian Cells. J Proteome Res. 2016 -03-04;15(3):1103-13.

56. Zheng Y, Thomas PM, Kelleher NL. Measurement of acetylation turnover at distinct lysines in human histones identifies long-lived acetylation sites. Nat Commun. 2013;4:2203.

57. Parthun MR. Histone acetyltransferase 1: more than just an enzyme? Biochim Biophys Acta. 2012;1819(3-4):256–63.

58. Cho NH, Cheveralls KC, Brunner A, Kim K, Michaelis AC, Raghavan P, et al. OpenCell: Endogenous tagging for the cartography of human cellular organization. Science. 2022 -03-11;375(6585):eabi6983.

59. Vaid R, Wen J, Mannervik M. Release of promoter-proximal paused Pol II in response to histone deacetylase inhibition. Nucleic Acids Res. 2020 -05-21;48(9):4877-90.

60. Ballas N, Battaglioli E, Atouf F, Andres ME, Chenoweth J, Anderson ME, et al. Regulation of neuronal traits by a novel transcriptional complex. Neuron. 2001 -08-16;31(3):353-65.

61. Martin BJE, Brind’Amour J, Kuzmin A, Jensen KN, Liu ZC, Lorincz M, et al. Transcription shapes genome-wide histone acetylation patterns. Nat Commun. 2021 -01-11;12(1):210.

62. Gryder BE, Pomella S, Sayers C, Wu XS, Song Y, Chiarella AM, et al. Histone hyperacetylation disrupts core gene regulatory architecture in rhabdomyosarcoma. Nat Genet. 2019 -12;51(12):1714-22.

63. Métivier R, Penot G, Hübner MR, Reid G, Brand H, Kos M, et al. Estrogen receptor-alpha directs ordered, cyclical, and combinatorial recruitment of cofactors on a natural target promoter. Cell. 2003 -12-12;115(6):751-63.

