## Supplemental Information for "Rapid degradation of Histone Deacetylase 1 (HDAC1) reveals essential roles in both gene repression and active transcription"

*To whom correspondence should be addressed

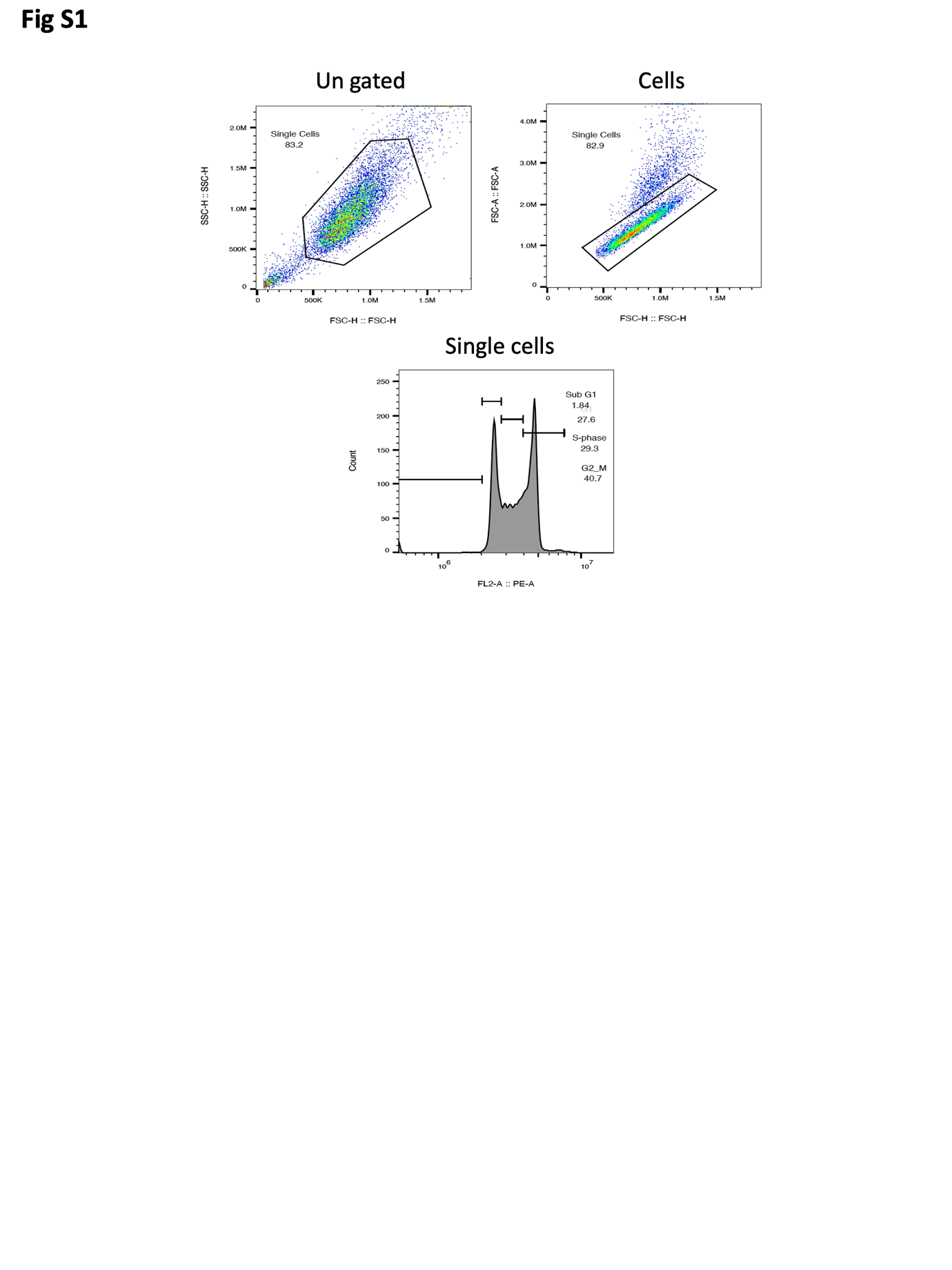

Figure S1. **Gating strategy used for PI FACS analysis.** A minimum of 10,000 events within the single cells gate were captured and analysed using FlowJo.

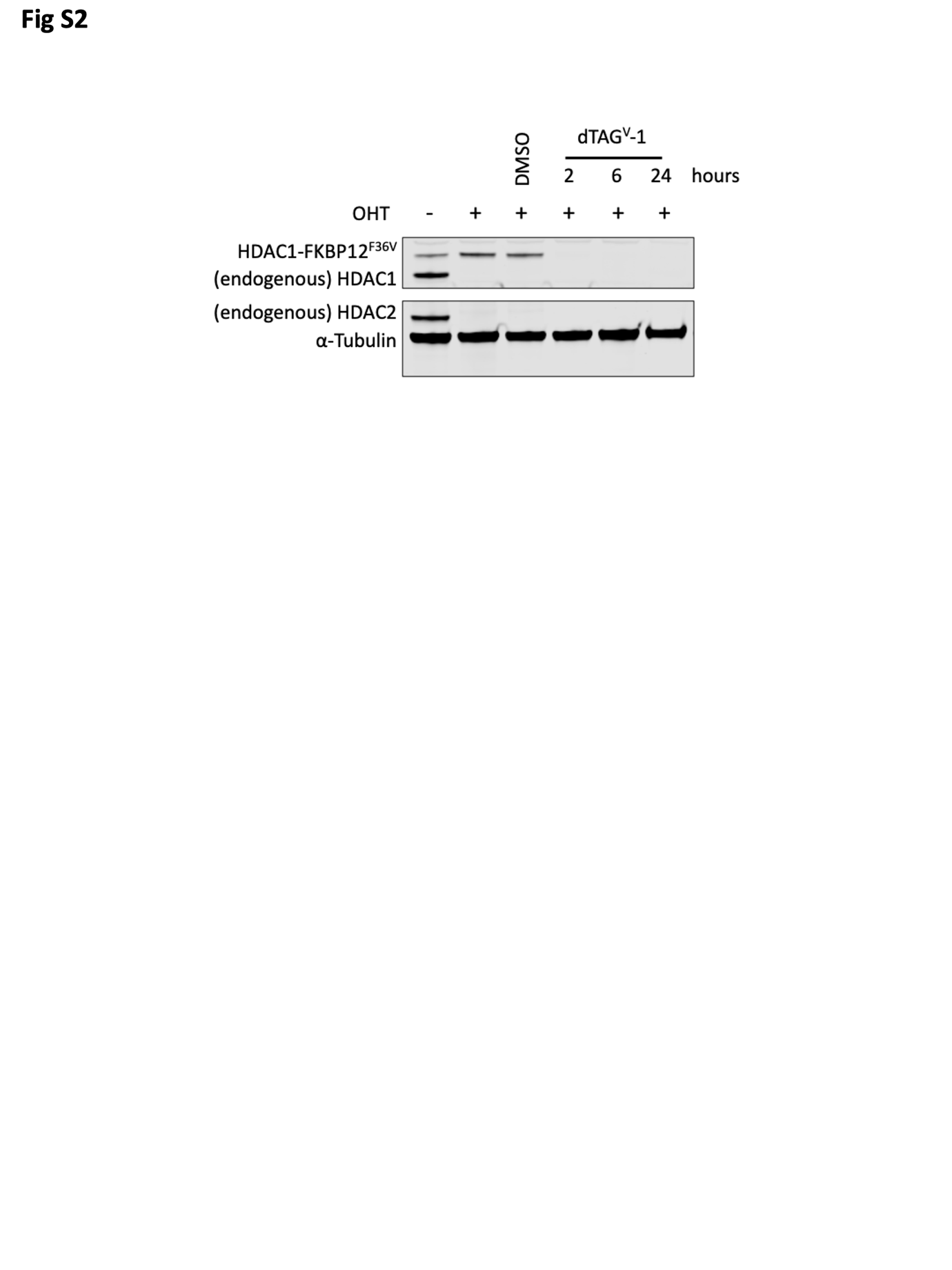

Figure S2. **dTAG^V^-1 degrades HDAC1-FKBP12^F36V^ efficiently.** Western blot showing endogenous HDAC1 and HDAC1-FKBP12^F36V^ (detected with ⍺-HDAC1), endogenous HDAC2 (detected with ⍺-HDAC2) when treated with OHT and dTAG^V^-1 as indicated, ⍺-tubulin shown as a loading control.

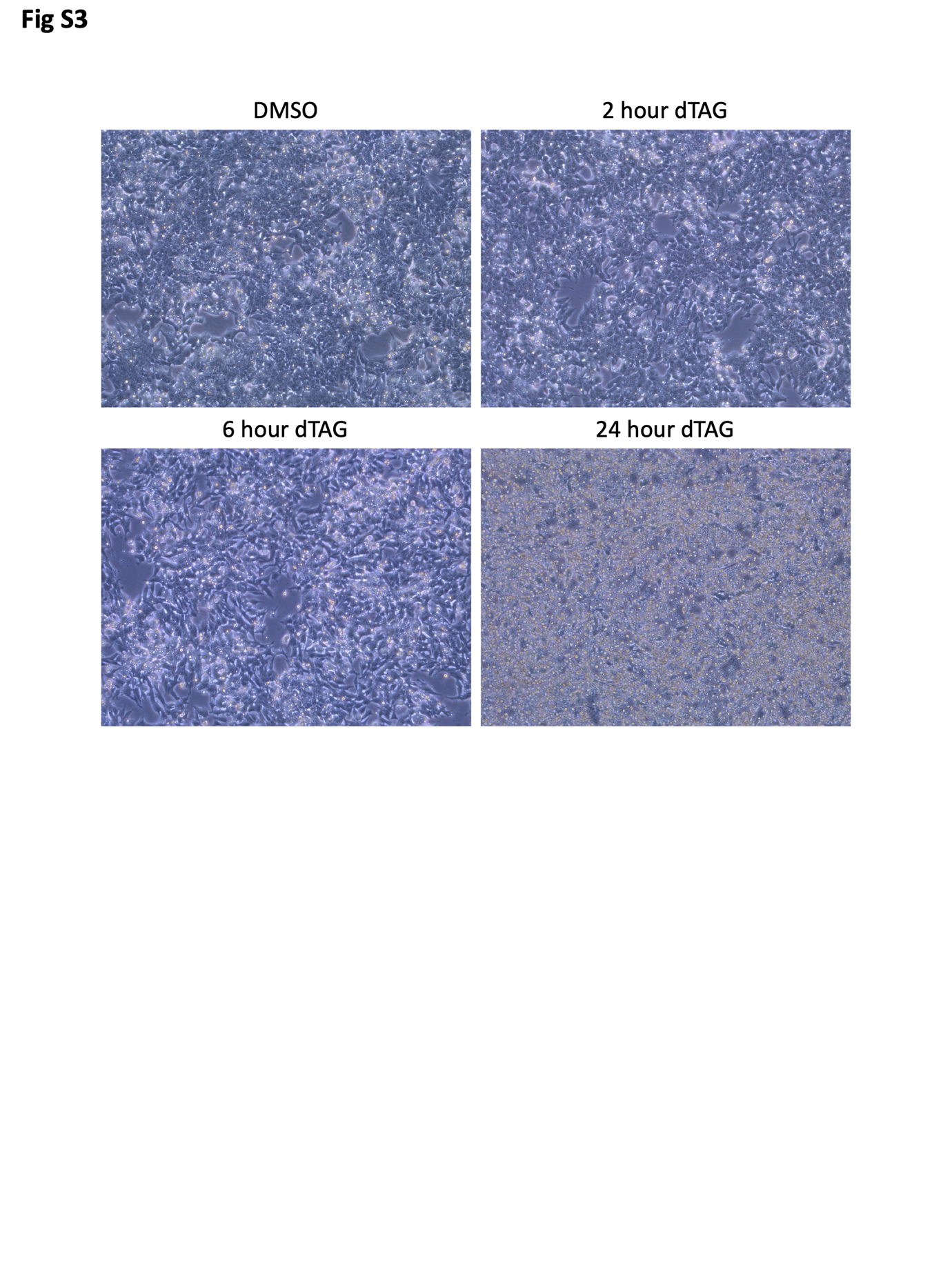

Figure S3. **HDAC1-FKBP12^F36V^ degradation causes cell death within 24 hours**. Images showing the effect of the indicated treatment time with 50 nM dTAG-13 on HDAC1-FKBP cells (images shown at 10x magnification).

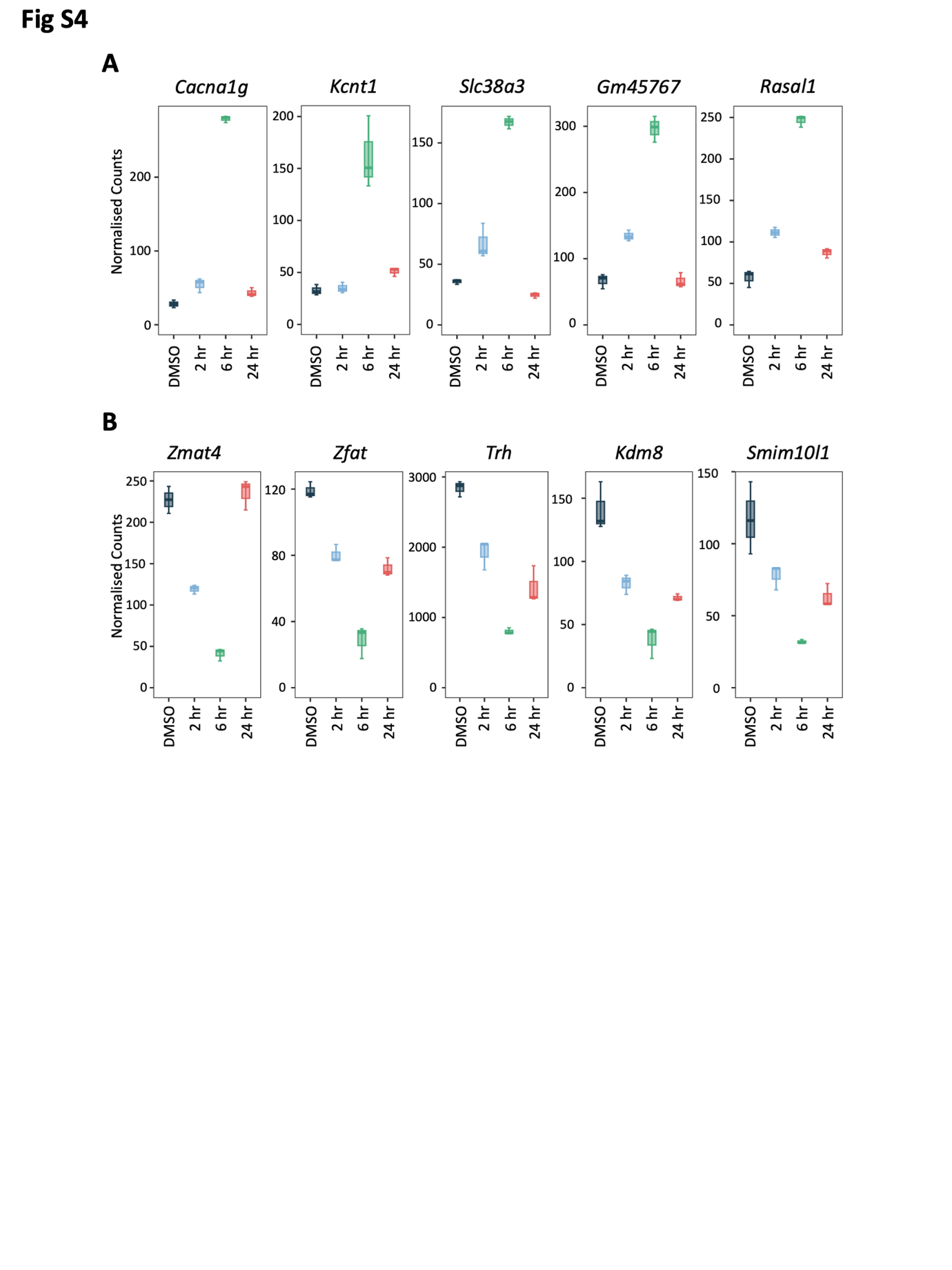

Figure S4. **A subset of genes is dysregulated only at 6 hours following HDAC1-FKBP12^F36V^ degradation.** (**A**) Boxplots showing the normalised count values for 5 genes which are only significantly upregulated (padj < 0.01, log_2_ fold change > 1) at 6 hours after HDAC1-FKBP12^F36V^ degradation. (**B**) Boxplots showing the normalised count values for 5 genes which are only significantly downregulated (padj < 0.01, log_2_ fold change > -1) at 6 hours after HDAC1-FKBP12^F36V^ degradation.

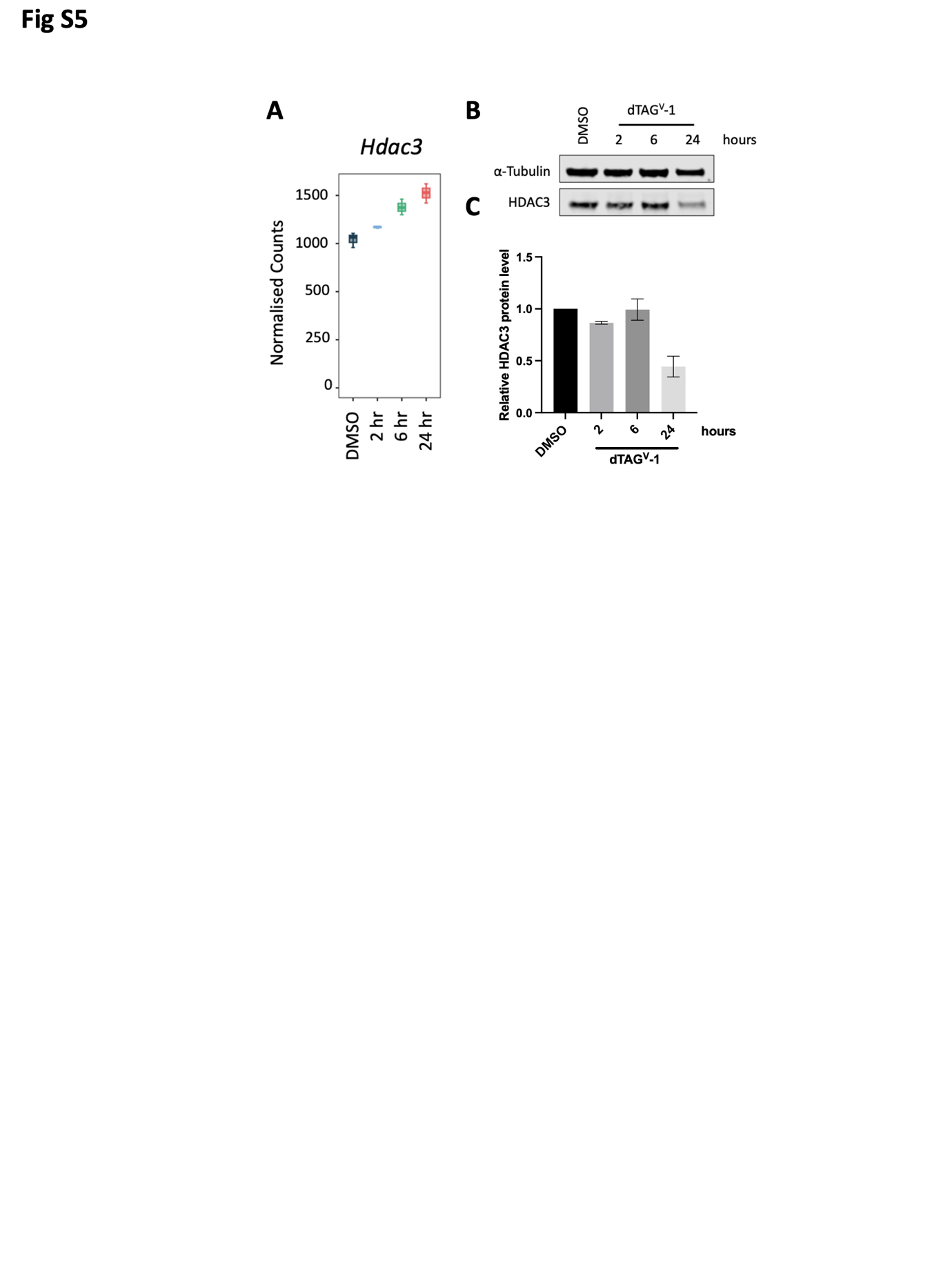

Figure S5**. There is not an increase in HDAC3 protein levels to compensate for loss of HDAC1/2**. (**A**) Boxplot showing normalised counts of *Hdac3* mRNA following the indicated dTAG^V^-1 treatment times. (**B**) Western blot showing HDAC3 proteins levels with the indicated dTAG^V^-1 treatment times, ⍺-tubulin shown as a loading control. (**C**) Quantification of blot shown on (B), bars show the relative levels of HDAC3 protein normalised to ⍺-tubulin (n=3, +/- SD).

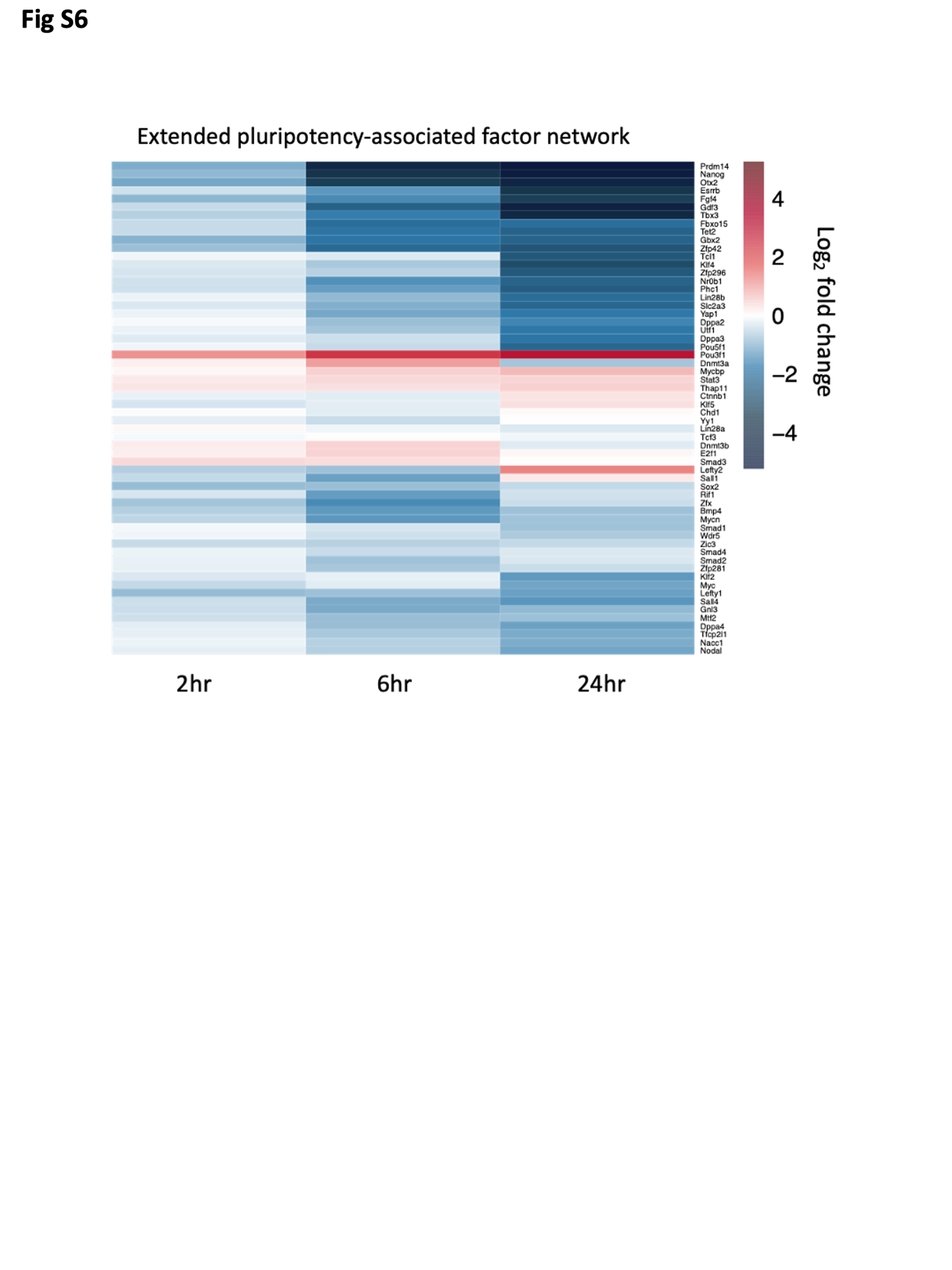

Figure S6. **HDAC1-FKBP12^F36V^ degradation causes a general downregulation of the extended pluripotency-associated factor network.** Heatmap indicating the log_2_ fold change values for the indicated pluripotency-associated genes at the indicated time points following dTAG^V^-1 treatment.

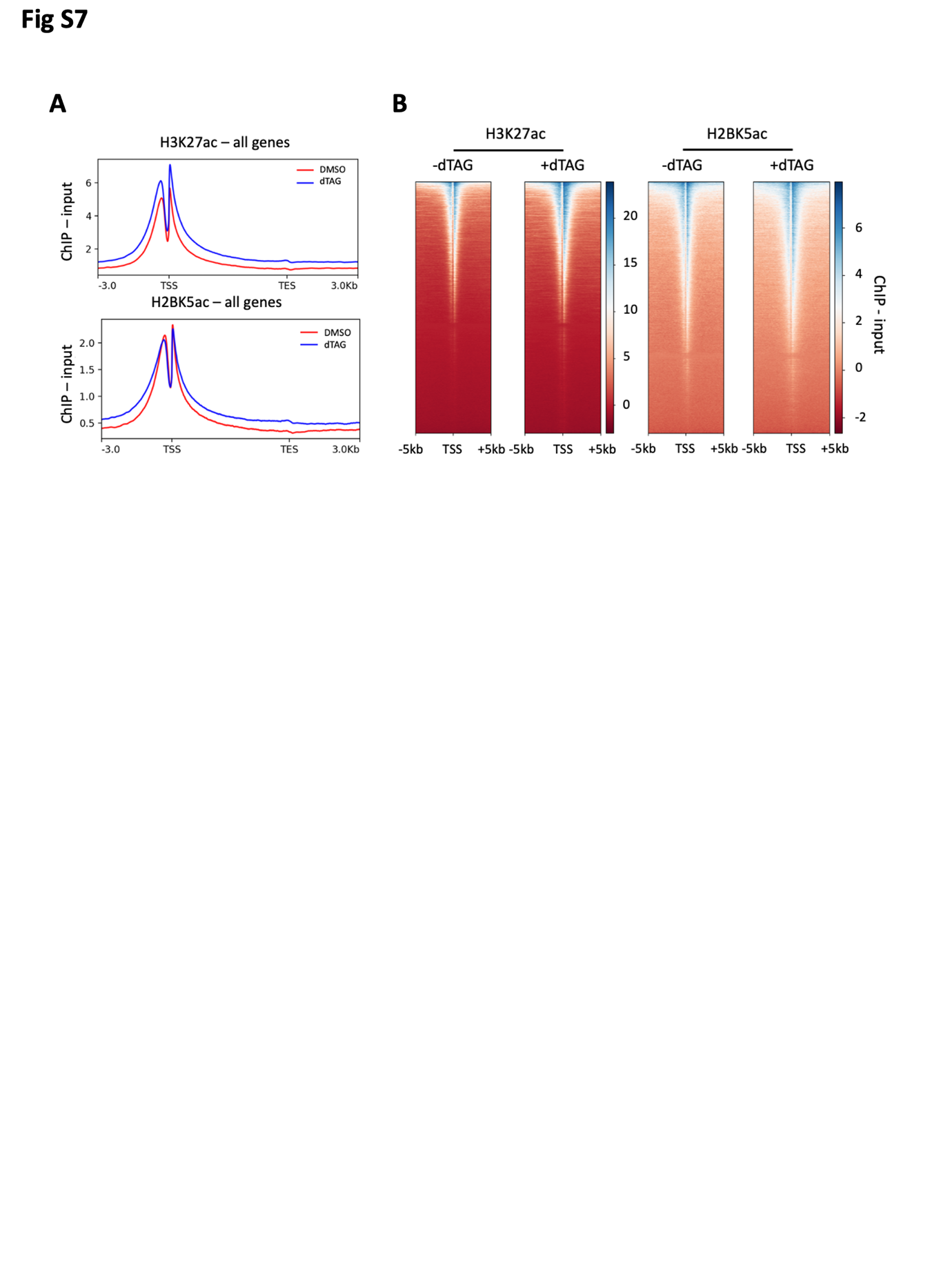

Figure S7. **H2BK5ac spreads due to HDAC1-FKBP12^F36V^ degradation.** (**A**) Metaplots showing the average signal (ChIP - Input) for H3K27ac and H2BK5ac across all genes following 6 hours of 100 nM dTAG^V^-1 or DMSO treatment as indicated, including the regions -/+ 3 kb from the transcription start site (TSS) and transcription end site (TES) respectively. (**B**) Heatmaps showing the ChIP - Input values for H3K27ac and H2BK5ac following 6 hours of 100 nM dTAG^V^-1 treatment as indicated, for the regions -/+ 5 kb from the TSS of all genes.

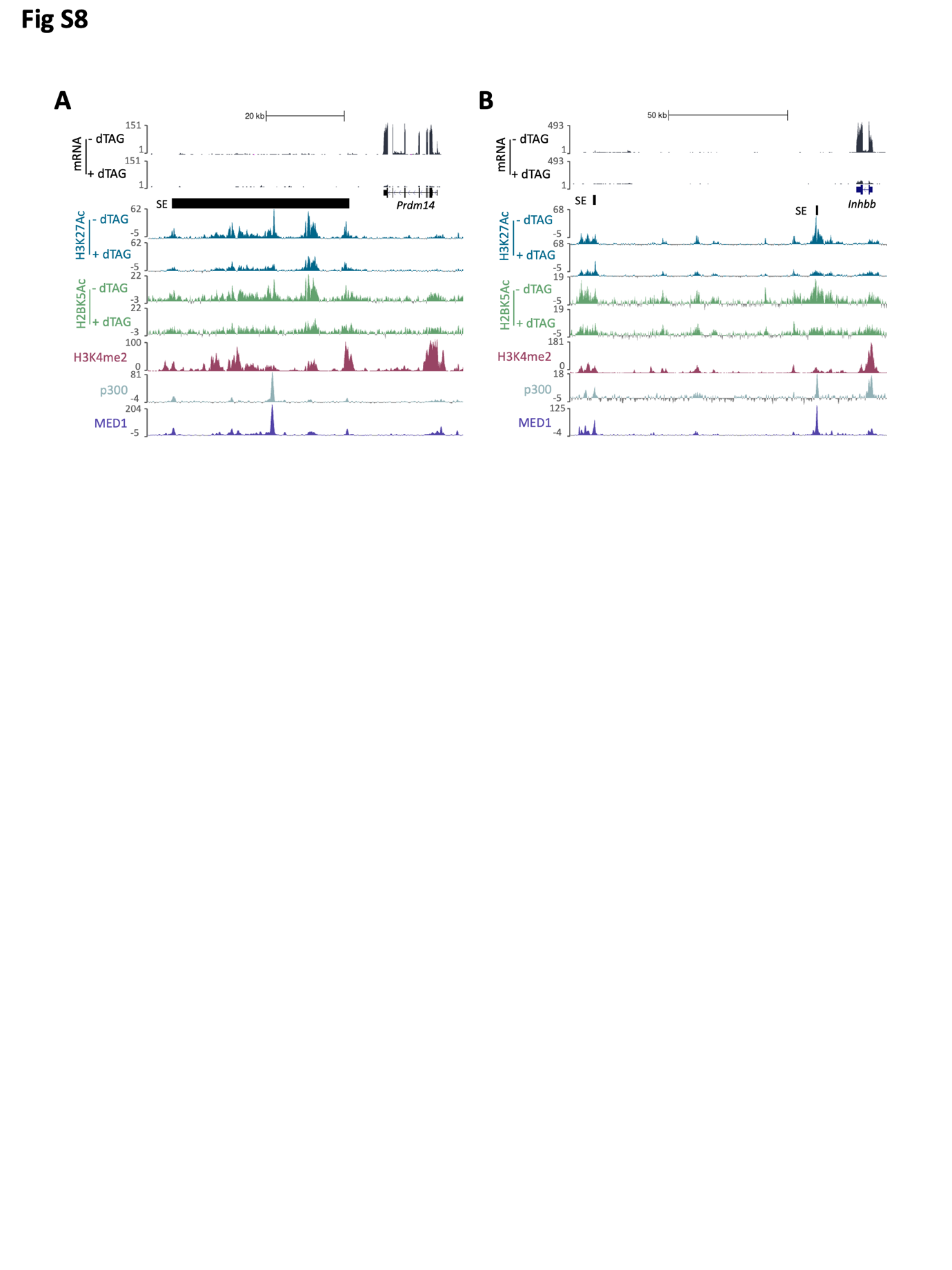

Figure S8. **The *Prdm14* and *Inhbb* super-enhancers show reduced H3K27ac and H2BK5ac which is commensurate with reduced gene expression.** (**A, B**) Tracks from the UCSC genome browser (37) showing the effect of 6 hours of 100 nM dTAG^V^-1 treatment on the mRNA, H3K27ac and H2BK5ac levels at the *Prdm14* (A) and *Inhbb* (B) loci. SE region is indicated. H3K4me2 (produced in our lab previously) and p300/MED1 (35, 36) ChIP data are also shown, highlighting the SE regions.

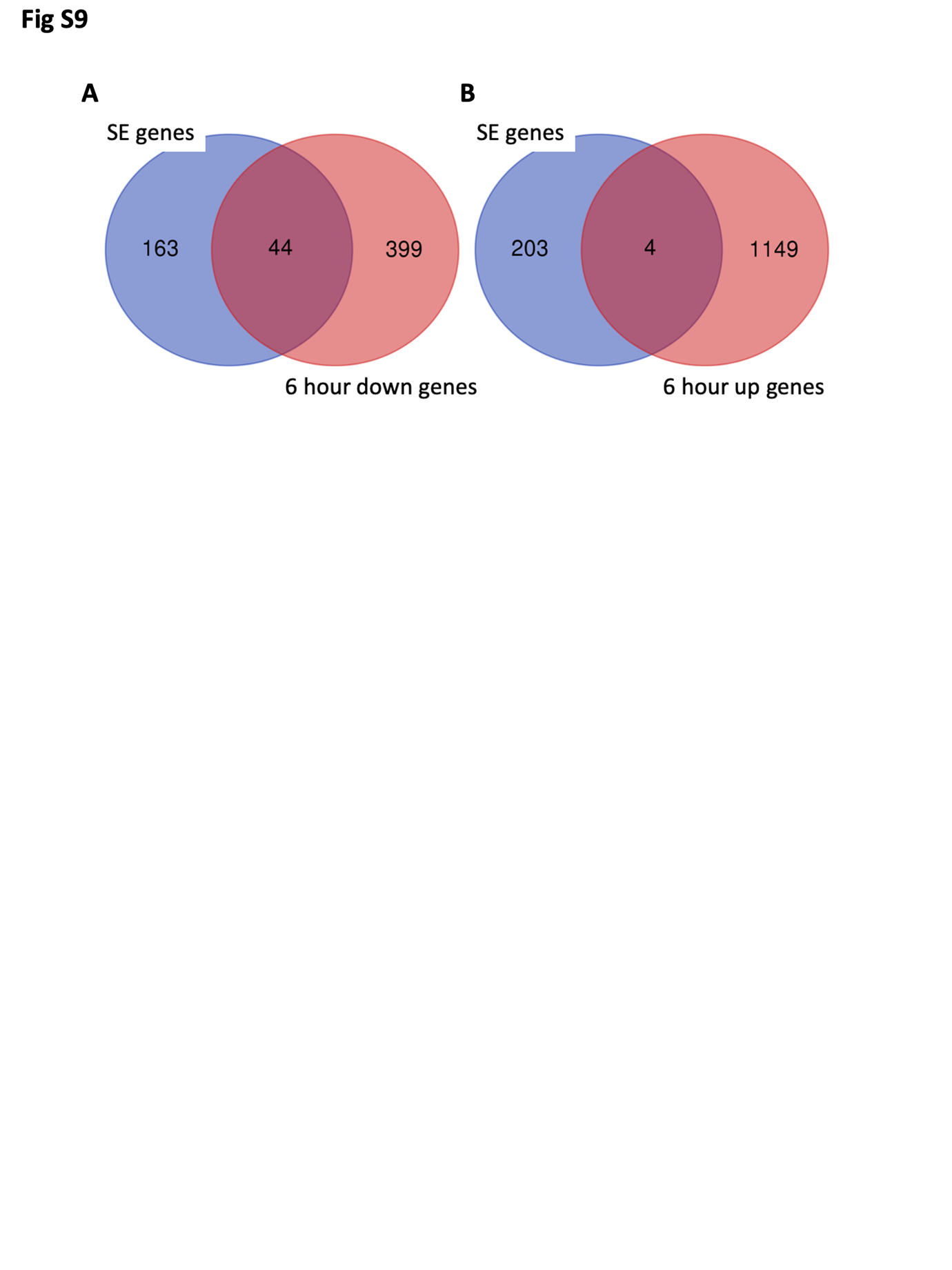

Figure S9. **Reduced super-enhancer (SE) acetylation is specifically linked to downregulation of gene expression.** (**A, B**) Venn diagrams showing the overlap between the SE regulated genes that we have assigned and the downregulated (A) or upregulated (B) genes determined by RNA-seq following 6 hours of dTAG^V^-1 treatment.
